# Flexible modulation of neuronal population dynamics drives variable decision-making in C. elegans

**DOI:** 10.64898/2026.09.03.749061

**Authors:** Jalaja Madhusudhanan, Anton Parinov, Charles Fieseler, Manuel Zimmer

## Abstract

Behavior arises from the interplay between spontaneous brain dynamics and sensory-driven responses, yet how spontaneous neural activity shapes variability in decision-making remains unclear. We leverage the tractable *C. elegans* nervous system to address this question. During oxygen avoidance, we observe binary trial-to-trial variability in behavioral responses. Whole-brain calcium imaging reveals a brain-wide sensory-to-motor transformation in which sensory neurons faithfully encode the stimulus but do not predict choice. Instead, decision-related information is distributed across interneurons and motor neurons, encoded through a neuronal subspace. This decision-biasing state evolves slowly during the pre-stimulus period, resembling preparatory dynamics for spontaneous behavioral transitions but receives neuromodulatory contributions. Optogenetic manipulations reveal that only a subset of neurons within this distributed representation are causally connected to choice. This reveals a dissociation between broad information sharing and control via dedicated localized nodes. Thus, response variability arises from slowly evolving modulation of brain states rather than stochastic circuit noise.

## Introduction

Animals constantly explore their environments driven by intrinsic needs, e.g., to locate food, find mates, seek shelter, or avoid predators. A considerable fraction of behavior is therefore spontaneously generated or occurs in an uninstructed manner, while the remainder arises in direct response to environmental stimuli. How intrinsically driven and sensory-driven behaviors are coordinated at the neuronal level remains poorly understood ^1^.

Intrinsic neural activity is a ubiquitous feature of nervous systems across species. Although once largely dismissed as background noise, spontaneous brain activity is now recognized as highly structured, reflecting ongoing internal computations rather than random fluctuations ^2,3^. Understanding the functional significance of these intrinsic dynamics has therefore become an outstanding question in neuroscience. Recent work across diverse model organisms has shown that a large fraction of ongoing neuronal dynamics encodes the animal’s current behavior, including both uninstructed and task-related actions. Strikingly, these behavioral representations appear distributed across the brain, extending even into primary sensory areas ^1,4–7^. Consequently, any sensory input must be integrated with these ongoing activity dynamics to generate or modify an appropriate behavioral response.

Perceptual decision-making typically involves a behavioral choice arising from a sensorimotor transformation process ^8–10^, and flexibility in this process gives rise to response variability. In mammals, decision-making research has primarily focused on sensory discrimination, learning and memory, the encoding of decision variables, and motor planning and execution ^11^. These studies have typically linked specific task parameters to dedicated brain regions such as the prefrontal cortex, premotor, and primary motor cortical areas where sensory evidence accumulation and motor preparatory activity can be localized ^8,9,12–16^. These findings are consistent with a longstanding framework in which specific cognitive functions are anatomically localized to dedicated brain regions ^17,18^. However, this view is increasingly challenged by large-scale neuronal recordings revealing that task-related information beyond motor commands is widely distributed across the brain ^19–22^. Moreover, individual neurons typically do not encode single task parameters in isolation but instead exhibit mixed selectivity, combining multiple variables within their responses ^23^. As a result, task-relevant information is thought to be encoded at the population level through neuronal subspaces, which are specific linear combinations of neural activity patterns that capture distinct computational variables within the same neuronal population ^23–25^. Yet the functions and mechanisms underlying brain-wide information sharing remain poorly understood. In particular, it is unclear whether all brain regions carrying information about a given task variable, such as a decision, are causally involved in decision-making or instead reflect corollary or downstream processing.

The distributed nature of brain computations raises the possibility of multiple sources of behavioral variability. Such variability could originate at any stage of sensorimotor transformation. It may arise at the sensory periphery, as in primate visual pathways ^26^ or at the level of motor execution, as in the fish cerebellum ^27^. It can also be introduced by dedicated variability-generating circuits, such as songbird cortical–basal ganglia pathways ^28^. Mechanistically, variability may stem from noise in individual circuit elements, such as synaptic variability ^29^ as demonstrated in *Aplysia* motor circuits ^30^. It may also reflect the instantaneous behavioral state, as during *C. elegans* turning ^31^ or initial neural population states, as in the premotor cortex ^10^. Finally, longer-lasting internal states can also bias decision-making, such as aggression-related hypothalamic states in mice ^32^.

Here, we leverage the tractable nervous system of the nematode *C. elegans* to comprehensively address these questions and elucidate the mechanisms of variable decision-making in the context of brain-wide intrinsic dynamics. *C. elegans* possesses a compact, stereotyped nervous system of only 302 neurons, comprising approximately 118 defined cell types with fully mapped connectivity ^33–37^. Its behavioral repertoire is organized around a core action sequence of forward crawling, backward crawling (reversal), and turning ^38^. Transitions between these actions occur stochastically even in the absence of acute sensory stimuli ^39,40^, but can also be triggered by sensory inputs, including fluctuations in ambient oxygen (O₂) concentrations detected by defined sensory circuits ^41^. Sensory modulation of behavioral transitions serves important ethological functions: *C. elegans* prefers intermediate oxygen concentrations of 5–12% and avoids atmospheric levels of approximately 21% ^42^. A shift from intermediate to atmospheric oxygen typically elicits a reversal, interpretable as an avoidance response ^41,43,44^. The transition from forward to backward crawling is typically preceded by a transient slowing or halting of forward locomotion ^4,40,45^ which may serve as a preparatory behavior enabling the abrupt change in crawling gait and direction ^4^.

Single-cell-resolution whole-brain recordings in *C. elegans* have revealed brain-wide population dynamics that reliably encode the animal’s action sequence ^4,46,47^. Here, multiple identified cell types are coupled to larger neuronal ensembles that show reproducible activation patterns tied to specific motor actions on every single event. For instance, the RIB interneurons, representatives for the forward motor command state, are selectively active during forward crawling, with activity levels correlating with forward locomotion speed ^4,48^. Conversely, the descending command interneurons AVA, representatives for the reversal motor command state, activate precisely at reversal onset and are inactive at the transition to forward locomotion ^4,49^. *C. elegans* therefore provides an ideal system in which to track sensorimotor transformation and variable decision-making both holistically and with single-cell resolution.

In this study, we designed a paradigm to study *C. elegans* behavioral responses to high-oxygen (21% O₂) and observed binary trial-to-trial variability in reversal responses. Whole-brain neuronal activity recordings reveal a brain-wide sensory-to-motor transformation flow and recapitulate this trial-to-trial variability in the reversal command state. Using this paradigm, we show that sensory circuits recruit the same brain-wide neuronal population dynamics observed during spontaneous motor command states. Sensory neurons, however, reliably report the stimulus and do not carry information about choice outcome, which instead can be best decoded from a distributed neuronal population state involving interneurons and motor neurons. This state evolves slowly during the pre-stimulus forward motor command period. It resembles the preparatory neuronal subspace preceding spontaneous reversal commands yet is distinct in its reliance on contributions from sensory-recruited neuromodulatory neurons. Using optogenetics and behavioral experiments, we find that only a subset of the neurons contributing to the decision-encoding state can be causally linked to behavioral choice. In summary, variability in decision-making in this system arises from a seemingly deterministic, slowly evolving brain state rather than from random stochastic fluctuations in circuit activity. Our data suggest that neuronal subspaces can be flexibly modulated to serve multiple functions. Moreover, we reveal a discrepancy between the representational structure of brain activity, whereby decision-related information is broadly shared across large neuronal populations, and more localized control knobs that can be causally linked to decision-making.

## Results

### Bimodal behavioral response variability to repetitive high-oxygen stimuli

To study the flexibility of behavioral responses of animals to identical sensory stimuli, we first established an experimental paradigm where we can reliably stimulate in a repeated fashion and read-out an array of behavioral responses (Fig. 1A). Previous studies on oxygen sensation in *C. elegans* have shown that their responses to fluctuating O_2_ concentrations is influenced by various factors like genetic background, feeding state, food availability, and other modalities ^41,44,50^. Therefore, we used *npr-1(ad609)* mutant worms in this study, which in previous studies served as a proxy for behavioral responses of wild *C. elegans* strains with elevated sensitivity to O_2_. Typically, these animals prefer intermediate O_2_ concentrations found in natural soil habitats or social feeding aggregates, and avoid atmospheric O_2_ levels ^42–44,51,52^. Moreover, we optimized the experimental conditions (e.g., temporal stimulus profile, feeding state, and food concentration) to match as many conditions as possible for behavioral recording and subsequent neuronal imaging (see methods). We performed high-resolution behavioral imaging of individual freely moving worms responding to changes in oxygen concentrations. Our stimulus protocol consisted of a 450-second baseline at 11% O_2_, followed by 16 trials of high-oxygen stimuli alternating between preferred 11% O_2_ (60 seconds) and aversive atmospheric concentrations of 21% O_2_ (30 seconds) (Fig. 1A, Fig. S1A).

**Figure 1:**
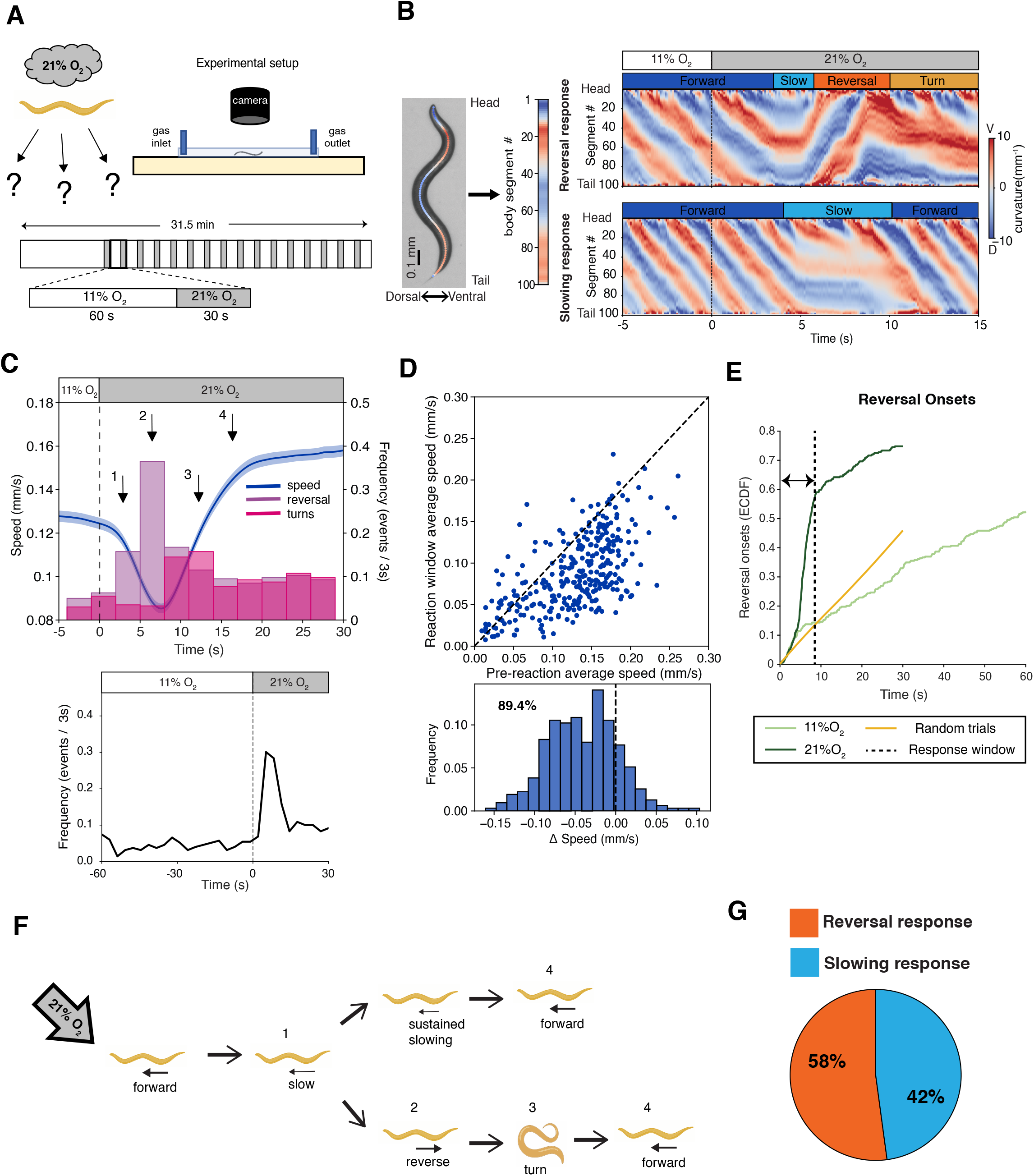
Worms show trial-to-trial response variability to repeated 21% O2 stimuli. **A.** Schematic illustrating experimental setup for behavioral recordings (upper) and stimulus protocol (lower). Each replicate consists of 16 stimulus trials. **B.** Worm image with skeleton overlaid (left) and corresponding curvature from head (segment 1) to tail (segment 100)(middle). The right panels show exemplary kymograms of posture time series. Bars show O2 concentration and current behavioral state. **C.** (above) Average forward speed (shading indicates SEM across all trials) and histograms of reversals and turns onsets. Arrows indicate the execution order of the behavioral sequence in response to 21% O2. (below) Stimulus-triggered reversal onset probability in 3-second bins. **D.** Upper: scatter plot of average forward speed from all trials, in a pre-reaction window (0-3 seconds) on the X-axis vs in the reaction window (4-8.5 seconds) on the Y-axis. Dotted black line (y=x). Lower: probability histogram of speed change (Y-X), fraction of slowing indicated. **E.** Empirical cumulative distribution function (ECDF) of reversal initiations over time into indicated stimulus episodes, across all trials. Light green, 11% O2 episodes, dark green, 21% O2 episodes. Random trials (orange) represent a shuffle trigger control (see methods). The dotted black line indicates the response window (0-8.5 seconds). **F.** Illustration of the two alternative action sequences executed by worms in the response window, numbered as in (C). **G.** Fraction of trial outcomes during response window. Data from N=25 replicates (different animals) / n=322 trials.

For each worm, we calculated posture kymograms, which capture most of C. elegans’ locomotion patterns and from which various secondary behavioral metrics can be derived (Fig. 1B). Firstly, we examined the responses to stimuli by annotating the major action sequence of worms, composed of forward crawling, backward crawling (reversal), and turns ^4,42^. Consistent with previous studies, in our experimental conditions, animals responded to oxygen upshifts by transient slowing of forward crawling speed, followed by upregulating reversals and turns, followed by accelerated forward crawling (Fig. 1C, Fig. S1B) ^43,44^.

Nearly all animals responded with initial slowing (∼89.4 % of all trials) (Fig. 1D), indicating reliable stimulus detection. Interestingly, we observed two further behavioral response categories (Fig. 1E-F). In more than half of all trials, animals initiated reversal/turn maneuvers within 8.5 s of stimulus onset (Fig. 1C, E, S1C-E). The remainder exhibited sustained slowing before accelerating forward crawling speed (Fig. 1G, 2A). To better distinguish between these two categories, we examined the response profile of reversal events across all trials. Reversals were upregulated within a few seconds of stimulus onset and returned to baseline after 8.5 s (Fig. 1E, S1C, D), defining a response window during which stimulus-evoked reversals can be statistically distinguished from spontaneous reversals, which occur at constant rates during baseline and 11% O_2_ periods. Based on this criterion, we categorized the response patterns into ‘reversal response’ (58% of all trials) versus ‘slowing response’ (42% of all trials) (Fig. 1G). Trial outcome structure from individual worms was not bimodal and rather revealed a spectrum of response probabilities, with many worms being representative of the entire population (Fig. S1E). This observation indicates that behavioral variability was not explained by persistent differences of individual worms during these experiments. However, we found that each trial outcome was not fully independent of response history: the probability of reversal outcomes was slightly elevated when the previous trial was a reversal response as well (Fig. S1F).

Our analyses further enabled us to compare evoked versus spontaneous actions. Both types were largely indistinguishable (Fig. S2), with comparable reversal durations, ventral versus dorsal turn biases, and turning strengths (Fig. S2B-G). However, reversal speed, pre-reversal slowing, and post-reversal acceleration were more pronounced in sensory-evoked events (Fig. S2A-C, E).

In summary, these behavioral data demonstrate substantial variability in the choice of actions when responding to each oxygen stimulus. While in nearly all trials animals slow down, demonstrating that the stimulus was reliably perceived, in only slightly more than half of the trials they recruit the major action sequence by upregulating reversal events. Variability arose on a trial-to-trial basis, with a persistent bias lasting at most two consecutive trials, or up to 180 s.

### Pre-stimulus behavior can partially predict but is not causally linked to behavioral choice

Next, we investigated whether variability in the pre-stimulus behavioral state could be a determinant of the observed variability in the post-stimulus behavioral choice. Previous studies showed that some of the worm’s response decisions indeed can depend on its ongoing behavioral state ^53,54^. Therefore, we extracted several instantaneous behavioral features, like forward speed, angular speed, curvature, and forward run length, as well as foraging-related long-term behavioral states like roaming and dwelling ^55–58^ from the behavioral raw data. Next, we pooled each trial by outcome categories and tested whether each feature was different in the pre-stimulus episodes. Notably, unlike other features, the forward speed contained some information about future choice: it was significantly lower in the reversal response trials compared to the slowing response trials (Fig. 2A). To quantify these observations for each of the individual behavioral features, we performed Linear Discriminant Analysis (LDA) ^59^, a supervised machine learning method, to estimate the prediction accuracy of each feature (Fig. 2B, C). This analysis showed that only locomotion speed could predict the upcoming behavioral choice, albeit with moderate improvement over the chance level, assessed via a shuffle control (Fig. 2A-C). Next, we tested whether a combination of behavioral features could improve the prediction accuracy. Therefore, we used a Partial Least Squares (PLS) regression analysis ^60^, a supervised machine-learning approach able to derive predictions from a combination of these behavioral features. For comparability with single feature models, we combined PLS with LDA (PLS-LDA) and calculated its cross-validated prediction accuracy. This model did not improve in comparison with our best single feature (forward speed) model (Fig. 2C).

**Figure 2:**
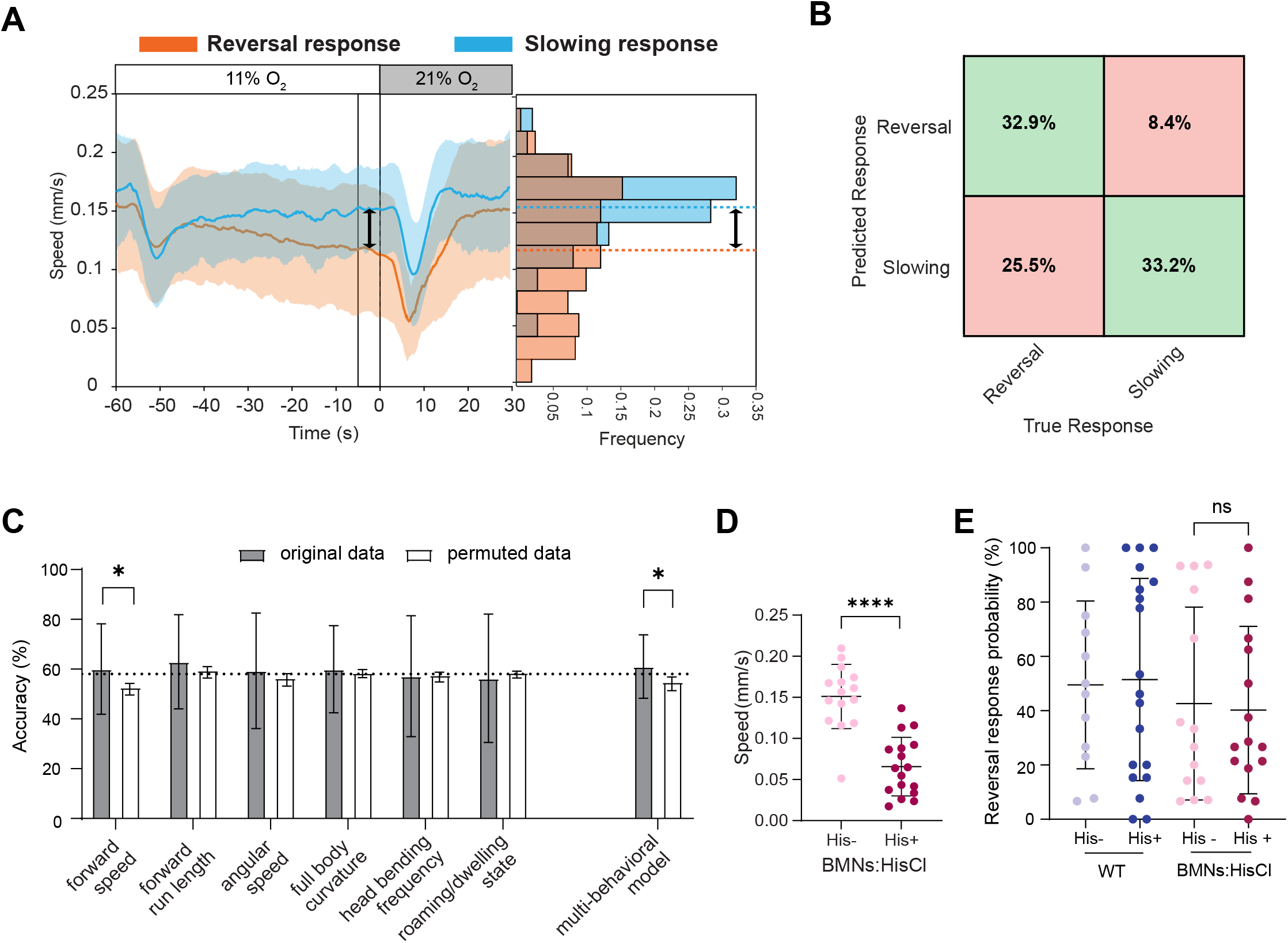
Forward speed can predict but cannot influence the response decision. **A.** Stimulus-triggered average forward speed in trials from two response categories (trial mean± std). The black line indicates the pre-stimulus time window (5 seconds) used to calculate the average speed values for the histogram on the right, and dotted lines show means. **B.** LDA confusion matrix showing the prediction accuracy of pre-stimulus forward speed as a predictor. Green = correctly predicted, Red = incorrectly predicted trials. **C.** LDA accuracies of individual behavioral features and the multi-feature PLS behavioral model with actual data (Gray, cross-validated mean±std) and shuffled data (White, mean±std). The dotted line indicates the chance level prediction accuracy (∼58%). * significantly higher than the shuffle control (see methods) (n=322 trials/ N=25 animals). **D.** Mean forward speed of animals with chemogenetic B-motor neurons inhibition (BMNs:HisCl), (Mann-Whitney test p-value ****<0.0001). **E.** Mean reversal response probability of wild type (WT) and BMNs:HisCl worms. Each datapoint in (D-E) represents a different animal, error bars show std (N = 12 animals, Mann-Whitney test).

Having identified locomotion speed as the sole, though weak, predictor of response outcome, we tested again whether this could be the result of a persistent property of each worm. Therefore, we compared the baseline speed of each individual with its response variability. However, we found no linear relation between these parameters, indicating that response outcome is not a consequence of individual differences in locomotion speed (Fig. S1G).

How could locomotion speed affect behavioral choice to reverse or not in response to an upcoming O_2_ stimulus? One hypothesis could be a biomechanical bias: since a switch from forward to backward locomotion typically includes an initial slowing maneuver ^4^ or pause state ^40^, starting from a slower locomotion speed might thereby favor future reversal initiations. This scenario would include a feedback mechanism reporting instantaneous locomotion speed to the decision-making circuitry. On the other hand, variation of locomotion speed could be a parameter controlled by a process further upstream of the motor periphery, which in parallel biases behavioral choice; in this case, pre-stimulus behavior would not be causally linked with decision-making. To distinguish between these possibilities, we targeted the B-Motor Neurons (BMNs) class, which are selectively required for the execution of forward locomotion in worms ^61^. Consistently, chemogenetic inhibition using a histamine-gated chloride channel (HisCl) ^62^ led to a reduction in the forward speed (Fig. 2D) without affecting the animals’ ability to execute reversals (Fig. 2E). However, this manipulation did not modulate response outcome (Fig. 2E). Interestingly, pre-stimulus locomotion speed, although much slower than in control animals, still separated between future behavioral choice (Fig. S3), suggesting that absolute speed is not causally related to the response decision. This result indicates that decision-making in this paradigm likely occurs via a process at an upper neuronal level controlling both locomotion speed and decision bias in parallel, motivating us in the present study to identify and characterize the corresponding neuronal state.

### A whole-brain activity map of sensorimotor transformation

To trace the process of sensory-to-motor transformation and to identify the potential sources of choice variability, we performed single cell resolution whole-brain calcium imaging, using a pan-neuronal nuclear localized Ca^2+^-indicator (nls-GCaMP6f) ^63^. Worms were immobilized in a 2-layer microfluidic device for controlled delivery of oxygen stimuli ^41,64^. We used worms with a pan-muscular expression of HisCl to inhibit muscles, hence transiently immobilizing the worms upon histamine treatment ^62,65^. We recorded the neuronal Ca^2+^ activities from N = 9 worms, applying a stimulus profile matched to the behavioral experiments above. Based on previously reported heuristics ^4,63,66,67^ (see also methods), we identified the cell type of 82 neurons. The number of datasets in which a neuron type could be identified varied between one to all nine animals (Fig. S4A, B). Fig. 3A shows a heatmap of a representative recording.

**Figure 3:**
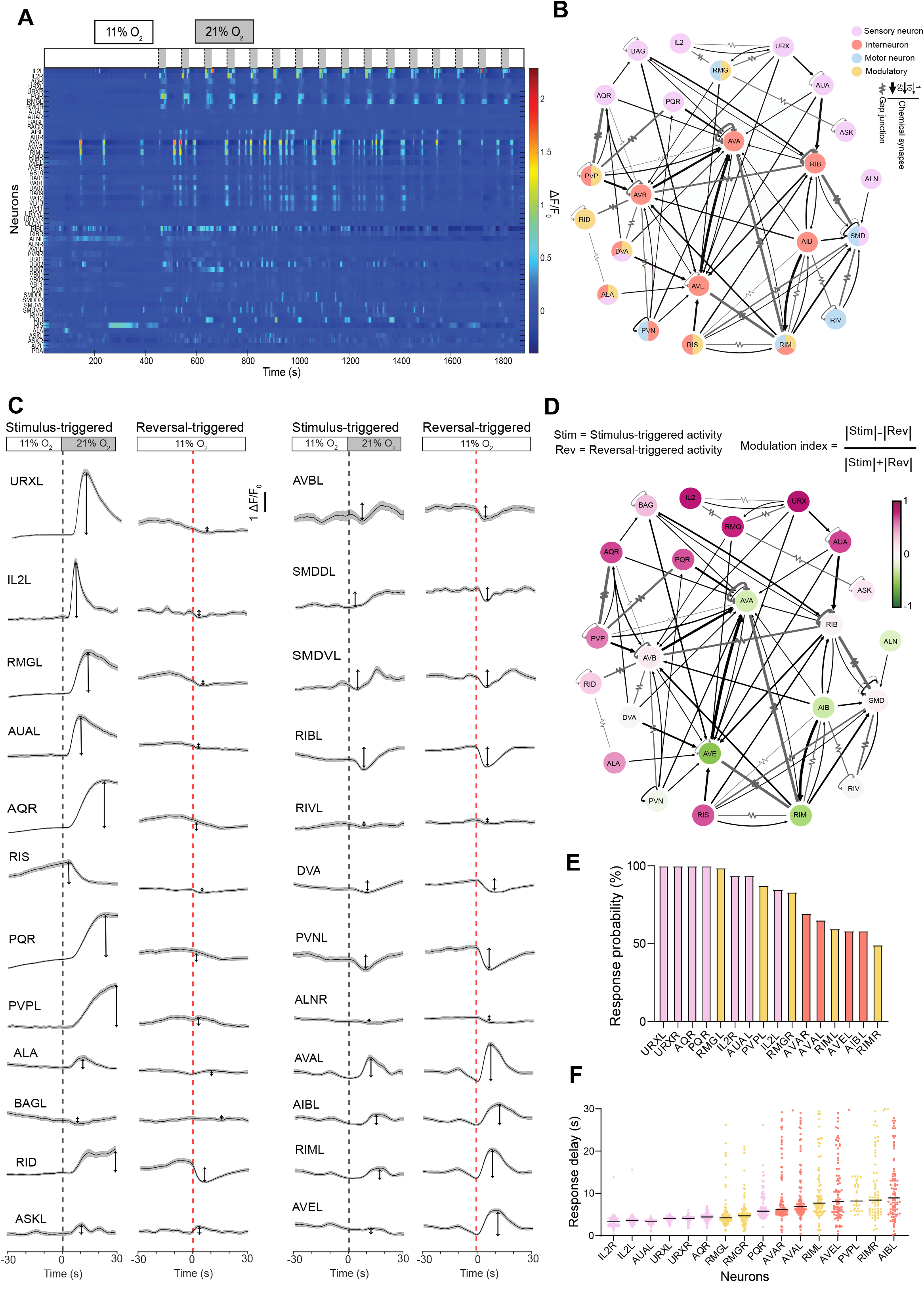
Whole-brain neuronal Ca2+ recordings capture sensorimotor dynamics. **A.** Heatmap of the neuronal traces (ΔF/F0) from an example whole-brain Ca2+ recording. Each row is a single neuron. **B.** Simplified oxygen chemosensory network created with the active neurons from our recordings. Modified from the available connectome resources (Nemanode.org). Colors indicate anticipated neuronal type (legend). **C.** Example trial averages of neuronal activity traces triggered to sensory stimulus (21% O2) or triggered to spontaneous reversal onsets (mean±SEM), black and red dotted lines show the trigger point. Arrows show the min-max difference in a 30-second window from the trigger point, indicating response magnitude. **D.** Overlay of the modulation indices (see formula) calculated from stimulus-triggered (Stim) and reversal-triggered (Rev) activities of each neuron (examples shown in 3C) on the oxygen chemosensory network in 3B. **E.** Ca2+-transient onset probabilities of indicated neurons within the 0-8.5s response window. **F.** Delay time to the first Ca2+-transient response of indicated neurons in all observed trials. Each dot represents a trial. Colors in (E-F) are consistent with (3B).

Previous studies showed that neuronal dynamics of *C. elegans* contain nervous system-wide coordinated activity patterns that reflect motor commands for initiating forward-crawling, reversals, or turning behaviors ^4,46,47,63,68^. Importantly, these dynamics are also apparent and largely preserved when animals are immobilized, enabling researchers to decode fictive behaviors ^4,47,63^. Specifically, the pre-motor neuron class AVA exhibits salient Ca^2+^-transients (Fig. 3A; Fig.4A), which in freely moving worms precisely align with the onset of reversal behaviors with single-trial reliability ^4,69,70^. We hence used these signals (“AVA onsets”) as a readout for the reversal motor command. Strikingly, the frequency of AVA onsets mimicked the pattern of reversal onsets in our behavioral experiments: upon 21% O_2_ stimulation, AVA onset frequency transiently rose from a baseline within 0-10 s (Fig. S4C, D). These data show that our experimental Ca^2+^-imaging paradigm faithfully recapitulates aspects of behavior observed in freely moving animals.

**Figure 4:**
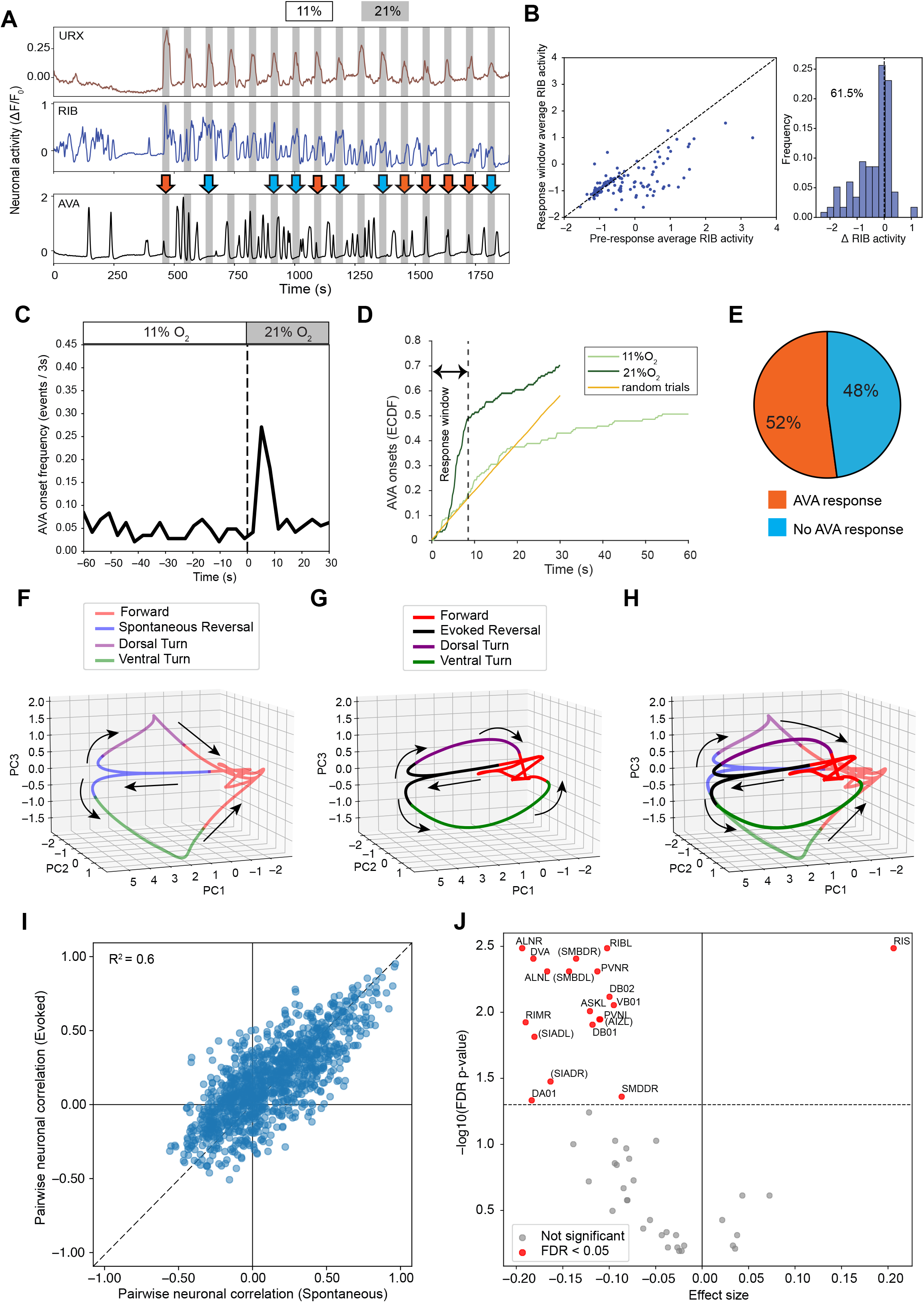
Immobilized recordings recapitulate response variability, and sensory-evoked reversal dynamics are comparable to the spontaneous reversal dynamics A. Example traces of URX (primary sensory neuron), RIB (forward interneuron), and AVA (reversal interneuron). Orange and Cyan arrows indicate, respectively, trials with AVA activity raising or not raising in the 21% O2 response window. B. (Left) Scatter plot of the average RIB activity from all trials, in a pre-reversal window (0-1 seconds) on the X-axis vs in the reversal window (4-8.5 seconds) on the Y-axis. Dotted black line (y=x). (Right) Probability histogram of the delta RIB activity (Y-X). C. Stimulus-triggered AVA Ca2+-rise onset probability across all trials. D. Empirical cumulative distribution function (ECDF) of AVA onsets over time into respective stimulus episodes, across all trials. Light green, 11% O2 episodes, dark green, 21% O2 episodes. Random trials (orange) represent a shuffle trigger control (see methods). The dotted black line indicates the response window (0-8.5 seconds). E. Fraction of trial outcomes during response window. Data from N=9 replicates (different animals) / n=117 trials. F-H. 3D plots of trial-averaged PC projections triggered to spontaneous (from 11% O2 period). Variance explained: PC1= 28%, PC2=17.6%, PC3=6.1%. (F) and sensory-evoked (from 21% O2 period) (G) reversal transitions, (H) is a merge of F &G. I. Scatter plot of Pearson correlation coefficients between peri-event activity profiles of identified neurons during forward-to-reversal transitions, comparing spontaneous transitions (x-axis) with sensory-evoked transitions (y-axis). The dotted blue line indicates the unity line (y = x). J. Volcano plot showing the effect size and significance of sensory modulation of peri-event activity profiles in identified neurons. Effect size is defined as the mean of the difference between the per-animal mean evoked activity and spontaneous activity. (p-values from custom permutation test, multiple comparison correction with FDR). Neurons in parentheses () indicate that neuron ID not verified and uncertain. indicates neuron ID not verified.

Next, we aimed at mapping sensory-to-motor flow onto the connectome of *C. elegans,* ranging from known O_2_ sensory neurons via first-layer sensory interneurons to pre-motor interneurons (Fig. 3B). To reveal how each neuron’s activity is modulated by the sensory stimulus, we first calculated their stimulus-triggered averages. This analysis showed that the stimulus propagated to all layers in the sensory-to-motor network (Fig. 3C, left columns). A signal in a stimulus-triggered average could have two major causes: (i) it could be a primary or secondary sensory response evoked by the stimulus; (ii) in addition, it could reflect a neuron’s coupling to the stimulus-entrained motor command state, which in *C. elegans* is composed of strongly correlated and anticorrelated neuronal activity ensembles. See ref. ^71^ for a detailed discussion. To disentangle these two potential sources, we calculated, from all 11% O_2_ episodes, the AVA onset-triggered averages to reveal how each neuron’s activity is generally modulated by the network-wide motor command (Fig. 3C, right columns). This analysis shows, in agreement with previous work ^4,46,47,63,67^, that the motor command related activity is widespread throughout interneuron circuits and typically does not or only weakly modulates the baseline activity of sensory neurons (Fig. 3C). To quantitatively compare both metrics, we calculated a ’modulation index’, which is positive for neurons whose activity is solely affected by the external stimulus or exhibit an otherwise stronger signal magnitude in evoked versus spontaneous reversals, 0 for comparable signal magnitudes, and negative for neurons whose motor response would be suppressed during sensory stimulation (Fig. 3D). Overlaying this metric onto the connectome shows a map of sensory-to-motor flow (Fig. 3D). It captures a set of sensory neurons (IL2, URX, AQR, PQR) and directly connected interneurons (RMG, AUA, PVP) primarily modulated by external stimulation. Interestingly, two arousal-related neuromodulatory interneurons not directly connected to any characterized primary O_2_ sensory neuron (RIS, ALA) also showed strong stimulus modulation (Fig. 3C-D). Notably, interneurons implicated in forward locomotion (AVA, RIB) showed little modulation, and those implicated in backward locomotion showed weaker stimulus-triggered than motor-triggered response profiles (Fig. 3C-D).

The negative modulation index in reversal interneurons could have multiple non-exclusive reasons. First, a neuron’s activity profile could indeed be amplitude-modulated by sensory circuits in individual trials, a possibility investigated further below. Additionally, a variation in response probability and/or response timing could lead to a lower averaged response profile. We therefore quantified both the likelihood (Fig. 3E) and timing (Fig. 3F) of signals in the post-stimulus period (at 21% O_2_) of selected neurons. (Fig. 3E). While most primary sensory neurons showed reliable (nearly 100% probability) and precisely timed responses within 1-4.5 s after stimulus onset, interneurons showed substantial variability, reflected in both reduced response probability and timing. This effect was particularly evident for reversal interneurons (AVA, AVE, AIB, RIM) (Fig. 3E-F).

In summary, these data demonstrate a sensory-to-motor transformation process evident in our experimental conditions. Our data indicate a segregation between a peripheral layer of primary sensory neurons and sensory interneurons, reliably reporting the external O_2_ stimulus, and an upper-level pre-motor interneuron layer, reflecting variability in the probability and timing of a reversal motor command. Surprisingly, a neuromodulatory neuron, RIS, was also recruited by sensory circuits.

### Whole-brain imaging conditions recapitulate binary choice paradigm

Having observed successful sensory-to-motor flow during whole-brain imaging conditions, we next investigated which aspects of the behavioral choice paradigm can be reproduced in this setting. As above, we used AVA activity onsets as a readout of reversal commands. Additionally, we analyzed the activity of RIB neurons, which are active exclusively during forward locomotion and whose activity linearly correlates with locomotion speed ^4,48^ (Fig. 4A). When animals encountered the 21% O_2_ stimulus during the forward command state and during phases of high RIB activity, the activity of this neuron reliably declined (Fig. 4B). Only in those instances, where RIB activity is already relatively low at the time of stimulus onset, no further decrease could be observed, suggesting a floor effect (Fig. 4B). This observation shares striking similarity with our locomotion speed behavioral data (Fig. 1D), suggesting that RIB activity can serve as a top-down control signal for locomotion speed, even in an immobilized animal.

Next, we analyzed the pattern of AVA onsets and found also striking similarities to our behavioral data: AVA onset frequency rose sharply from a stable baseline within a window of 0-8.5 s after stimulus onset before returning to baseline (Fig. 4C-D, S4C-D). Hence, we could use the same response window of 0-8.5 seconds, as applied to the behavioral data above, to also classify two choice outcomes from our imaging data: 52% AVA response (equivalent to reversal response) trials, and 48% No AVA response (equivalent to slowing response) trials (Fig. 4E). Moreover, as in our behavioral data, choice probability across individual worms was a spectrum indicating that choice variability arose on a trial-by-trial basis (Fig. S4E). Also consistent with the behavior, we observed a weak inter-trial dependency, with AVA responses being more likely when the previous trial had an AVA response as well (Fig. S4F).

In summary, both processes, unrestrained O_2_-regulated behavior as well as the O_2_ sensory-to-motor transformation process observed in the Ca^2+^-imaging conditions, remarkably shared major features: both exhibit an initial slowing state, followed by a binary choice of initiating a reversal state versus not. Compare Fig. 1D with 4D, Fig. 1C, E with Fig. 4C, D, and Fig. S1 D-F-G, respectively, with Fig. S4D-F.

### Evoked responses mirror spontaneous neuronal population dynamics

As shown above and in previous work, *C. elegans* explores its environment through spontaneous behavioral transitions — such as forward-reversal switches — that occur in the absence of acute external stimuli ^4,40,47,63^. The internal neural dynamics underlying these transitions involve coordinated activity across a large population of interneurons and motor neurons, which can be captured as a low-dimensional manifold reflecting the stereotyped action sequence of forward locomotion, reversal, and turning ^4^. Since the sensory-evoked behavioral transitions largely mimicked spontaneous behaviors in our paradigm, we tested whether sensory circuits recruit distinct or identical neuronal activity patterns to control these behaviors. In the previous section, we developed a method to group brain-wide reversal commands into those evoked by sensory stimulus (0-8.5 s response window) vs spontaneous ones (all events outside the response window) (Fig. S5A-B). As previously, we used Principal Component Analysis (PCA) to calculate the low-dimensional manifold representation of the action sequence and compared both categories. This analysis revealed qualitatively the same neuronal state trajectories aligning in this latent space (Fig. 4F-H), indicating that the stimulus-evoked brain-wide dynamics largely resemble spontaneous action dynamics. To quantify this finding, we tested whether the underlying neuronal correlation structure is conserved across both categories. For this purpose, we calculated the pairwise neuronal correlations spanning forward-to reversal state transitions and compared both categories via a linear fit (x=y with R^2^=0.6), showing that both neuronal population states largely align (Fig. 4I). This is also evident in the reversal onset-triggered averages of individual neurons, when separated into evoked vs spontaneous events (Fig. S5C). However, in many of them we observed amplitude modulations (Fig. S5C), which are quantified in Fig. 4J. This effect is mostly found in interneurons and motor neurons implicated in forward locomotion, reflecting a stronger pre-reversal downmodulation in stimulus-evoked transitions (Fig.4J, S5C). Contrarily, the neuromodulatory neuron RIS showed pre-reversal upregulation in evoked events (Fig.4J, S5C), an observation we investigate further below.

In summary, these results show that O_2_ sensory circuits recruit qualitatively the same neuronal dynamics that can be observed during spontaneous events. However, consistent with enhanced slowing in behavioral experiments (Fig. S2B), the pre-reversal forward state termed ‘slowing’ ^4^ appears stronger downmodulated prior to transitions to evoked reversals.

### Ongoing interneuron dynamics correlate with future behavioral choice

What could be the origin of variability in our binary choice paradigm? Connectomics analyses suggested a layered sensory-to-motor flow architecture in the worm brain ^72,73^. Therefore, we next systematically investigated at which layer response variability arises in the current paradigm.

Our previous analyses showed that choice variability is evident at the level of the pre-motor interneuron AVA (Fig. 4C-E), which is the major connectivity output to the motor circuitry required for executing reversal behavior ^33^. These A-class motoneurons consistently responded to AVA (Fig. S5C). Hence, we can exclude the motor periphery executing reversals as a major origin of choice variability. In conclusion, we hypothesized that the response decision must originate at the level of sensory and/or interneurons. To investigate this, we grouped neuronal activities in each trial into the two outcome categories (Fig. 4E) and compared the stimulus-triggered averages of all neurons in these two categories (Fig. S6A). It should be noted that a difference in the post-stimulus episode, especially during the response window, should reflect the execution of the choice, i.e., the motor command itself, as is evident in reversal interneurons and A-class motor neurons (e.g., AIB, AVE, RIM, and VA01, DA07) (Fig. S6A). Notably, some head motor neurons (potential SIAs and SMBs) showed salient and reproducible activity profiles after stimulus onset in the slowing response category (Fig.S6A). This observation supports the classification of a binary choice paradigm, i.e. choice between two alternative responses as opposed to response vs no-response.

Unlike activity features characterizing the neuronal response of the animal to the stimulus, a difference in the pre-stimulus forward state or early during stimulus presentation indicates a neuronal activity correlate of the decision-making process. Interestingly, the primary oxygen sensory neurons (URX, AQR, PQR) showed no differences between both categories throughout each trial, suggesting that these neurons reliably report the stimulus and excluding their primary responses as sources of choice variability (Fig. 5A, S6A). Unlike in the activity profiles of sensory neurons, we found in multiple interneurons strikingly different activity profiles between both categories; Fig. 5B shows three prominent examples. Notably, the difference in these examples emerged in the pre-stimulus time up to 30s prior to the stimulus onset. To quantify these observations across all identified neurons systematically, we performed frame-by-frame T-tests (FDR corrected) throughout the peri-stimulus interval, which indicates those time-intervals where each neurons shows significant differences between both categories (Fig. 5C). This analysis revealed two sensory interneuron classes (PVPL and AUAR) and, strikingly, other interneuron classes (RIBL, RIBR, RIS, DVA, PVNR) with prolonged continuous episodes of significantly different activity, up to 30s pre-stimulus onset (Fig. 5C). We next complemented this frame-by-frame analysis with a method suitable for rigorous cross-validation. We therefore trained LDA classifiers for each individual neuron and found a subset with prediction accuracy above chance level and significantly higher than a shuffle control (Fig. S7A). Interestingly, the best of these models (RIB) already outperformed the behavioral models; however, prediction performance was modest (Fig. 2C, Fig. 6A). These data indicate that a neuronal population state during the pre-stimulus time could underlie behavioral choice variability.

**Figure 5:**
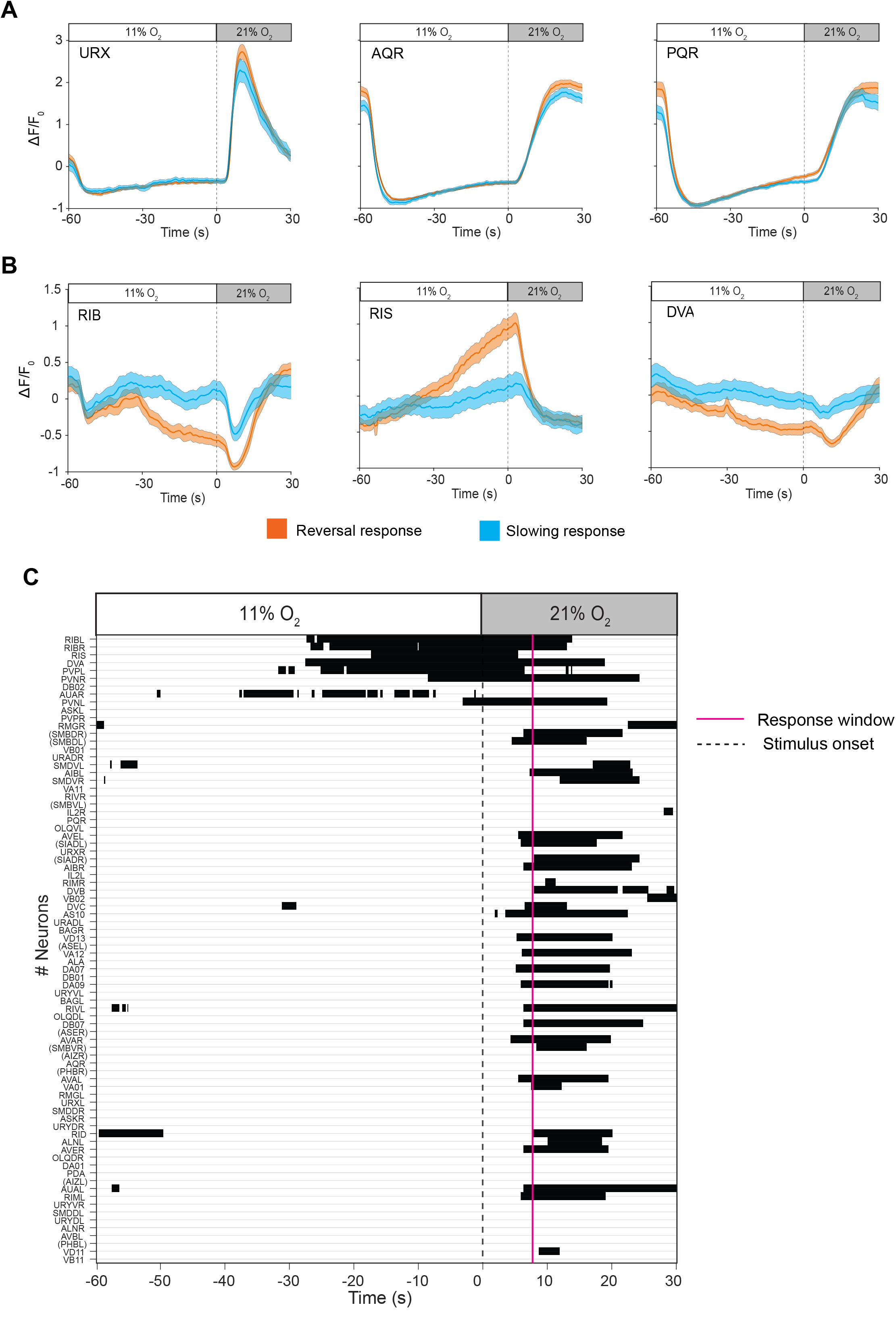
Neuronal explanation for response variability. Exemplary stimulus-triggered average traces of **A.** primary oxygen sensory neurons (URX, AQR, and PQR), and **B.** example interneurons (RIB, RIS, and DVA) in the two response categories. (trial mean±SEM) Orange = Reversal response trials, Cyan = Slowing response trials. **C.** Statistical comparison (T-test, with FDR correction for multiple comparisons) of stimulus-triggered traces between the response categories in individual neurons. Black=p-value<0.05, Neurons are sorted based on the lowest total p-value in a -20-second to +2-second pre-stimulus window. Neurons in parentheses () indicate that neuron ID not verified and uncertain.

**Figure 6:**
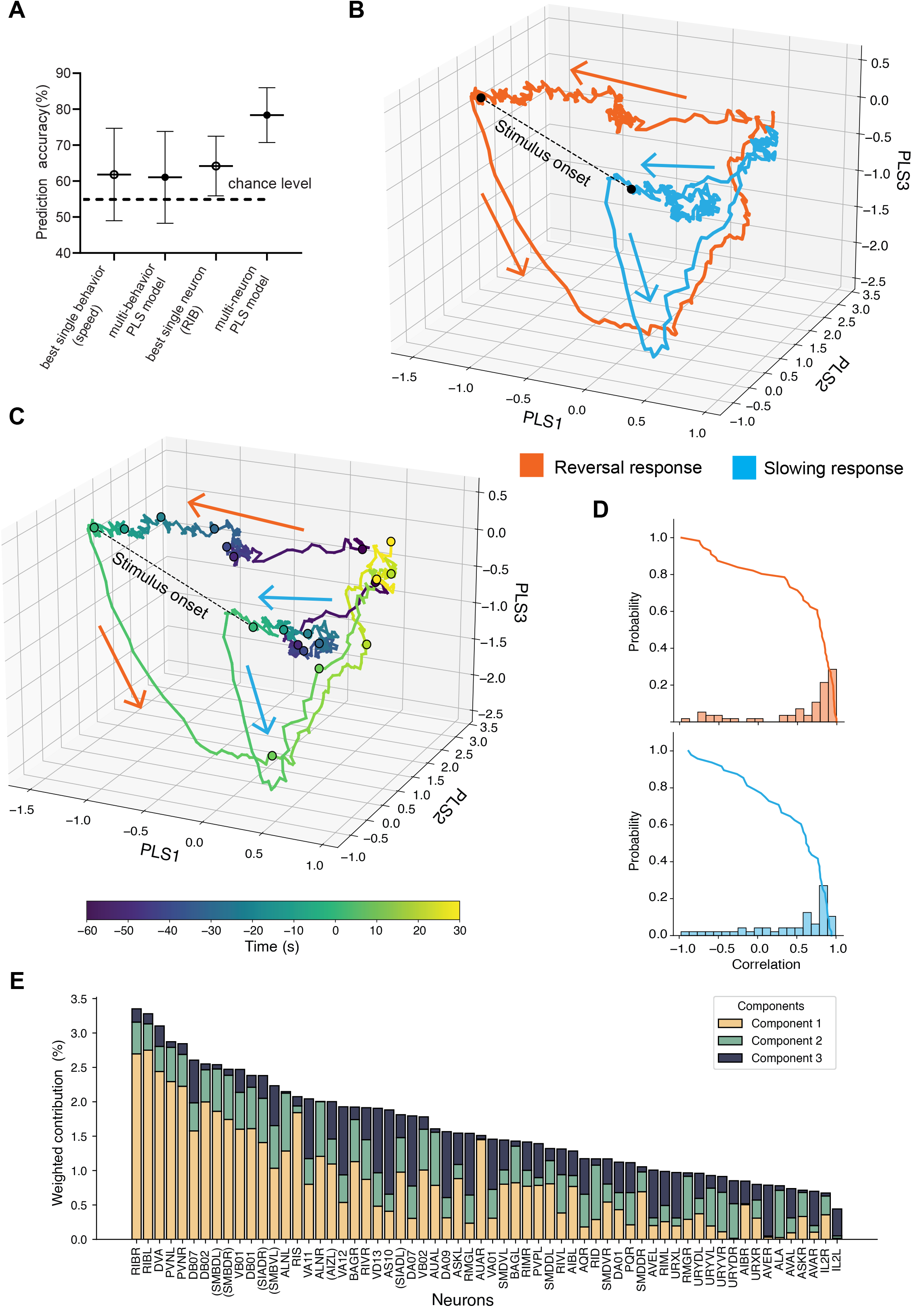
Population models show distributed pre-stimulus prediction states. **A.** Cross-validated LDA prediction accuracies of the best single feature and multi-feature PLS model from behavior (same as Fig. 2C) and neuronal recordings. The dotted line indicates the chance level prediction accuracy (∼55%). **B.** 3D plot of average PLS traces in the reversal (Orange, n=61trials/N=9 animals) and the slowing (Cyan, n=56 trials/N=9 animals) response trials. **C.** Same 3D plot of average PLS traces in B, color-coded by time. **D.** Histogram of correlation coefficient between individual trials and the average after PLS-LDA projection (see methods) in the reversal response (top) and slowing response (bottom) categories. The solid line shows complementary (rightward) cumulative probability. **E.** Weighted contribution (loadings) of each neuron to the top 3 PLS dimensions (components), indicating individual neurons’ contributions (feature importance) to the multi-neuronal PLS model. Feature importance of each component (PLS1 = 18.5±0.3%, PLS2 = 11.1±0.3%, and PLS3 = 4.2±0.2%) (see methods). Neurons in parentheses () indicate that neuron ID not verified and uncertain.

### Choice bias correlates with a distributed brain state operating on slow timescales

Our observations from individual neurons motivated us to ask whether a multi-neuron model could yield improved performance. PLS is particularly useful for studying neuronal population dynamics because it identifies a shared latent space by jointly decomposing neural activity and categorical labels, isolating the population-level structure most relevant to distinguishing between conditions (see methods). Therefore, we built a multi-neuronal PLS model from the pre-stimulus neuronal activity. To evaluate this model’s performance in a comparable way to the previous models, we used LDA and calculated the cross-validated prediction accuracy of the PLS model (PLS-LDA), yielding ∼80% (see methods). The neuronal population model outperformed the behavioral models (Fig. 2C), as well as the best single neuron model (Fig. 6A, S7A). To characterize population activity relevant to the behavioral choice, we projected the neuronal data into the PLS latent space and plotted the average trajectories of reversal and slowing response trials. Consistent with the single neuron observations, these trajectories drifted apart as early as 30 s prior to stimulus onset during the forward state (Fig. 6B, C). This appearance led us to hypothesize that a latent brain state evolving on a slow timescale underlies the animal’s bias to engage in one behavioral choice versus the other. Alternatively, such a brain state could rapidly switch in a more binary and stochastic fashion; hence, the slow drift could be a result of averaging. To distinguish between these possibilities, inspired by ref. ^74^, we investigated whether such progressive drift is seen also at the level of single trials. Despite obvious trial-to-trial variability, most individual instances indeed reflected such drifting behavior in their one-dimensional LDA projections (Fig. S8). We aimed to quantify these observations and thus calculated the correlation of individual trials with the average in each respective category. In both response categories, ∼ 80% of trials showed a positive correlation, indicating that the trial average indeed captures some common representative features (Fig. 6D). However, the presence of low and even negative correlations in individual trials (Fig. 6D), as well as fluctuations within the individual trial trajectories (Fig. S8), both indicate substantial single-trial heterogeneity as well.

Finally, we investigated the contributions of individual neurons to the population PLS model by calculating their weighted feature importances (see methods) (Fig. 6E). This analysis indicates which neuronal circuitry is potentially implicated in the choice bias and its underlying brain state. We found that interneurons (RIB, DVA, PVN) and motor neurons (DB07, DB02), typically active during forward locomotion, as well as head motor neurons implicated in turning (potential SMBs), as well as the neuromodulatory neuron RIS, ranked on top by their model contributions, while O_2_ sensory circuits (IL2, URX, AQR, PQR, RMG, AUA, PVP) contributed less (Fig. 6E). Importantly, the sorted spectrum of neuronal contributions fell off smoothly, lacking any obvious cutoff. This result indicates that information about the ongoing choice bias is not bounded within a small set of neurons but distributed across the brain.

In summary, our analyses support a model in which behavioral choice emerges from a decision bias encoded in a slowly evolving brain state that is broadly distributed across diverse cell types, including neuromodulatory neurons, interneurons, and motor neurons, with small effects from O_2_ sensory neurons. Conceptually, this is different from a brain state that operates hierarchically or orthogonally to motor circuitry; instead, both decision-bias and execution seem to be interwoven.

### Preparatory brain states preceding spontaneous and evoked behavioral decisions exhibit shared and distinct neuronal features

Given its intertwined nature, it is possible that the neuronal state reflecting choice bias corresponds to a preparatory state during which the system is generally prone to switch from forward to backward crawling. In this case, a small perturbation like a sensory input would accelerate the same transition underlying spontaneous events. Alternatively, and equally interesting, it could be a state specifically biasing the system’s responsiveness to the external stimulus. To further distinguish between these possibilities, we first visualized the average neuronal population trajectories leading to spontaneous and sensory-evoked reversals in the latent space of the PLS model (henceforth, ‘sensory PLS model’) (Fig. 7A). The corresponding trajectories took different paths, which is consistent with distinct neuronal subspaces. A 2D projection of the underlying data points within the forward state showed two distinct distributions; however, with substantial overlap (Fig. 7B).

**Figure 7:**
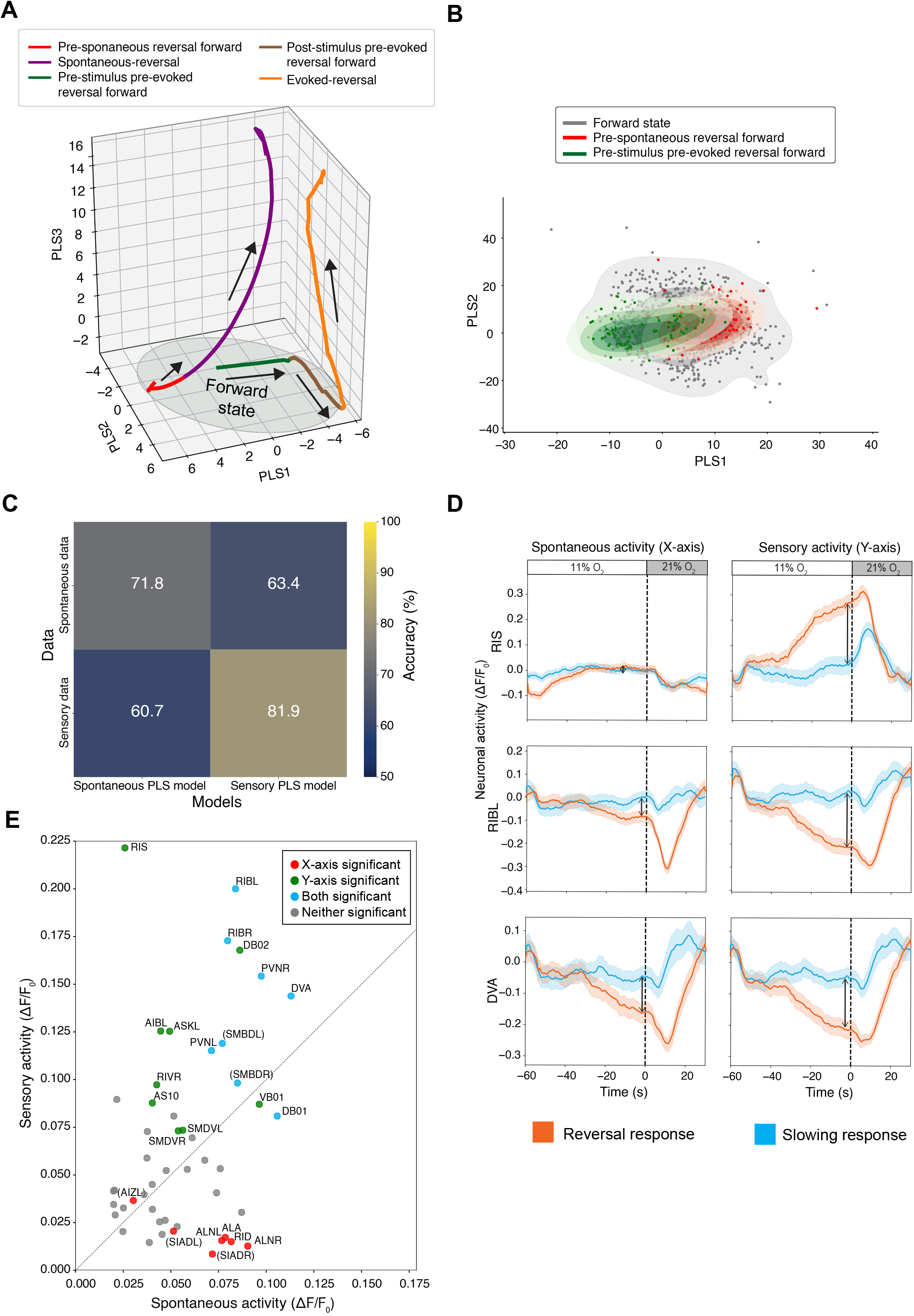
Sensory-evoked reversal decision originates from a distinct neuronal subpopulation. **A.** 3D plot of average PLS traces leading to sensory-evoked (Orange) and spontaneous reversals (Purple). Average traces of pre-stimulus, preparatory activities of evoked (Green) and spontaneous (Red) reversals are also shown. **B.** Scatter plots and corresponding 2D contour density plots of neuronal activity state during forward episodes, projected into the first two PLS dimensions. Each datapoint represents the average of PLS 1 and 2 dimensions for an 8.5-second time window leading to spontaneous reversals (Red), sensory-evoked reversals (Green), and the rest of the forward state (Gray). **C.** Confusion matrix showing accuracies of LDA models predicting the indicated state transitions, using either the sensory-reversal PLS model or the spontaneous-reversal PLS model (see methods). The chance level accuracies are 52.1% and 51.62%, respectively. **D.** Stimulus-triggered traces (mean±SEM) of example neurons in the reconstructed neuronal traces from the spontaneous PLS model (spontaneous activity), and in the reconstructed neuronal traces from the sensory PLS model (sensory activity) (see methods). The black arrow indicates the maximum difference between reversal and slowing response trials in a -20-second pre-stimulus window (effect sizes). **E.** Scatter plot showing effect sizes of neuronal modulation by choice category as predicted by spontaneous vs sensory PLS models. A high value on the x-axis indicates neurons whose choice modulation prior to stimulus onset is captured by the spontaneous PLS model, whereas a high value on the y-axis indicates neurons whose choice modulation prior to stimulus onset is captured by the sensory PLS model. Red = X-axis significant, Green = Y-axis significant, Cyan = Both axes significant, and Gray=Not significant. Statistical test: custom permutation test, p-value<0.05, multiple comparison correction with FDR. Neurons in parentheses () indicate that neuron ID not verified and uncertain.

To quantify any differences between these subspaces, we first trained another PLS model to classify upcoming spontaneous reversals out of the forward state (henceforth, ‘spontaneous PLS model’) (see methods). The weighted feature distributions of the sensory and spontaneous PLS models were largely similar, with only a few distinctions, e.g. the neuromodulary neuron RIS ranked relatively high or low, respectively (compare Fig. 6E versus Fig. S9). Next, we cross-validated the performance of these models using PLS-LDA (see methods). The resulting confusion matrix showed that both models performed well in predicting their corresponding upcoming state transitions (diagonal entries in Fig. 7C). Interestingly, both PLS models showed moderately good performance (>60% vs 50% chance level) in predicting the respective other state transitions (off-diagonal entries in Fig. 7C). In summary, even though the preparatory neuronal subspaces for spontaneous and sensory-evoked reversals have some shared features, they also use distinct information.

Next, we aimed to compare both subspaces at the neuronal level in an intuitive manner. For this, we reconstructed each neuron’s activity trace using both spontaneous and sensory PLS models and calculated the stimulus-triggered averages in both choice categories. Any divergence in the trajectories illustrates whether the respective model captures the neuron’s modulation by the behavioral choice. Fig. 7D shows selected examples of neurons that are modulated by choice outcome in the pre-stimulus episodes as shown in Fig. 5B. The activity of neuromodulatory neuron RIS diverges between the categories prior to stimulus onset (Fig. 5B, 7D). This difference is well captured by the sensory PLS model but not by the spontaneous PLS model (Fig. 7D). This indicates that RIS activity contains unique information about sensory choice bias. Interneurons RIB and DVA show modulations when reconstructed by either model. This indicates that RIB and DVA contribute to choice bias in a similar fashion as they contribute to upcoming spontaneous reversals. To explore these features systematically across individual neurons, we generated a neuron-wide scatter plot of the maximum trajectory differences obtained from spontaneous (x-axis) vs sensory (y-axis) model projections (Fig. 7E). We found that the neuromodulatory interneuron RIS stood out in its uniqueness, reflecting a neuronal state preceding sensory-evoked reversal responses. Furthermore, forward interneurons like RIB, DVA, and PVN reveal modulation in both models. Notably, most of these neurons’ modulations appeared well above the diagonal in Fig. 7E, indicating that the neuronal state encoding choice bias is generally more pronounced than the neuronal state predicting spontaneous reversals (See also Fig. 7D); this is also consistent with differences in the triggered averages of sensory-evoked versus spontaneous reversals (Fig. S5C). Other interneurons like RID, ALA, and SIADs reveal modulation only in the spontaneous model subspace (Fig. 7E); this reflects their general modulation when approaching reversal events, which, however, did not inform the sensory PLS model.

In summary, these analyses indicate two distinct but related preparatory neuronal subspaces carrying information about upcoming spontaneous and sensory-evoked reversals. We described a preparatory neuronal subspace in which the system is generally prone to switch behavioral state from forward crawling to reversal. Additional contributions from arousal-related neurons RIS and other interneurons, in particular RIB, can shift this state further, biasing the system’s responsiveness to the sensory stimulus.

### Pre-stimulus activity state of neurons RIB and RIS, but not DVA or B-MNs, can causally influence behavioral choice

Our analyses of neuronal population activity revealed a distributed brain state correlating with a decision bias and action selection. Finally, we aimed to test whether the thereby identified neuronal circuits can also causally influence the decision-making process. Therefore, we applied chemogenetic (via HisCl) and optogenetic (Chrimson) perturbations to a set of selected neurons identified in our models. As shown above, inhibiting B-MNs did not influence the decision bias (Fig. 2E), despite the strong effect on locomotion speed (Fig. 2D), and despite their major contribution to the sensory PLS model (Fig. 6E), and the preparatory state predicting spontaneous reversals (Fig. 7D). We next targeted several interneurons. Based on our analysis results, we focused on RIB interneurons implicated in forward locomotion speed ^4,48^, DVA neurons, a stretch receptor interneuron located in the tail implicated in body posture and explorative behaviors ^75–77^, and RIS neuromodulatory neurons implicated in sleep and locomotion speed ^45,66,78^. We validated the effectiveness of the perturbations by testing for their expected effects on baseline behavior, i.e., locomotion speed and body posture (Fig. S10A-C, H). To test the neuron’s causal implication in behavioral choice bias, we evaluated the effects of perturbations on behavioral choice upon O_2_ stimuli (Fig. 8 and Fig. S10E-F, I). Based on its activity patterns (Fig. 5B) and model contributions (Fig. 6E, 7D), we expected that RIB activity would counter the behavioral choice to reverse, while low RIB activity should promote the reversal choice. Confirming our expectations, chemogenetic inhibition of RIB neurons led to a strong bias towards higher reversal response outcomes (Fig. 8A). Note that RIB inhibition had a similar effect on locomotion speed as B-MN inhibition, which did not affect choice bias (Fig. S10A, Fig. 2D). Conversely, activating RIB acutely via optogenetics for 15 s before the 21% O_2_ stimulus (Fig. S10D) led to fewer reversal response decisions (Fig. 8D, Fig. S10E).

**Figure 8:**
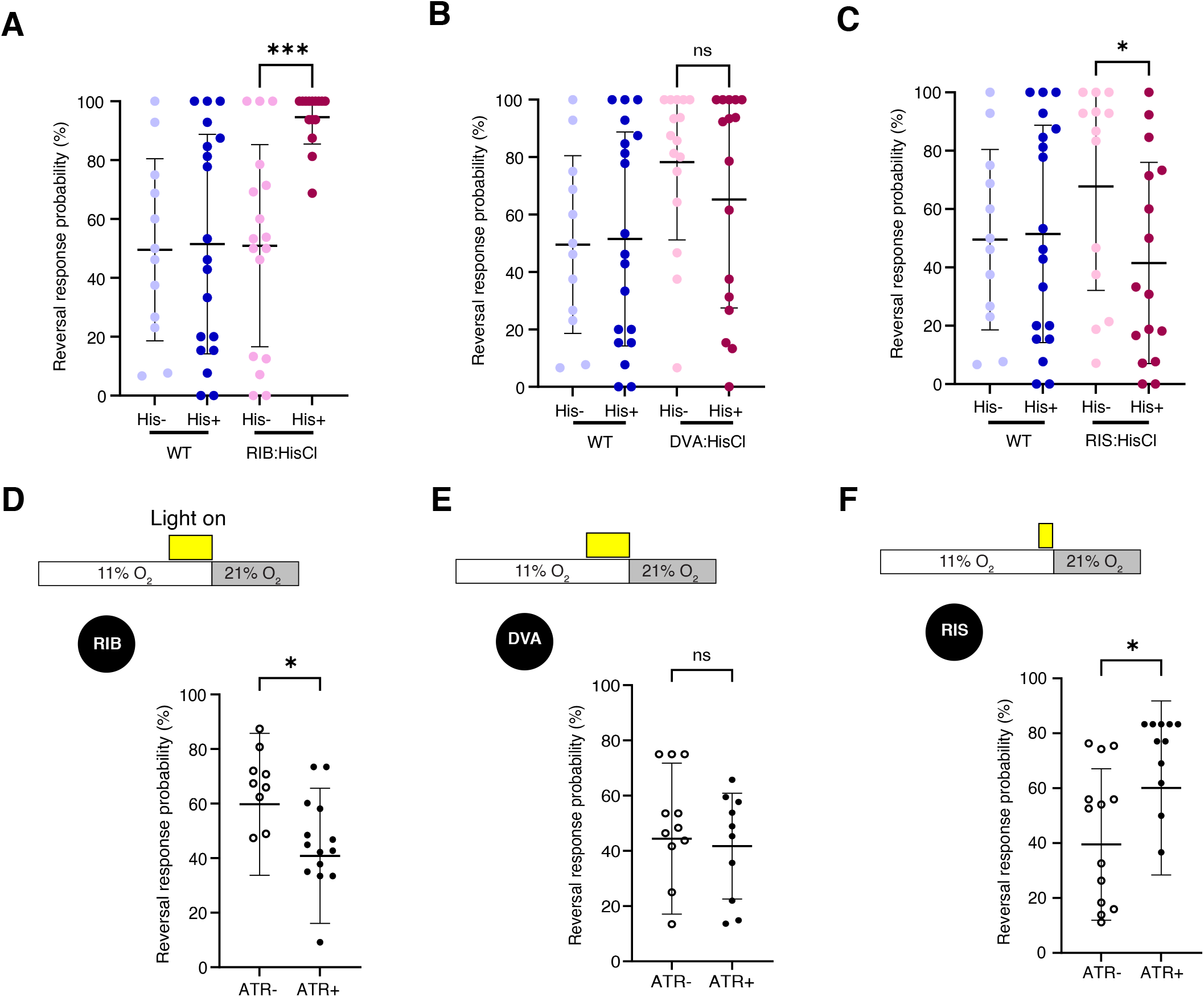
Pre-stimulus activities of RIB and RIS neurons can causally influence the response decision. A-C. Chemogenetic inhibition of interneuron RIB (**A**), DVA (**B**), and RIS (**C**). **D-F.** Optogenetic activation of interneuron RIB (**D**), DVA (**E**), and RIS (**F**). Bars indicate the timing of gas and light stimulus used in each trial for each neuron. For optogenetic experiments, the data were normalized using a Light-only control (see methods details and Fig. S10 E, F & I for the non-normalized data). Each datapoint represents an animal, error bar shows mean± std (N => 10 animals, Mann-Whitney test p-value ****<0.0001, ***<0.001, **<0.01, *<0.05). Negative data were not plotted, but used for mean calculations and statistics.

Based on its activity patterns (Fig. 5B) and model contributions (Fig. 6E, 7D), we expected that DVA activity, similar to RIB, would counter the behavioral choice to reverse, while low DVA activity might promote the reversal choice. However, both chemogenetic inhibition and optogenetic activation of DVA neuron did not alter the response variability (Fig. 8B, E, Fig. S10F), indicating that perturbing DVA activity before stimulus onset does not influence the behavioral choice.

In comparison to RIB, the RIS neuron showed a complementary activity pattern prior to the behavioral choices; typically, its activity rose prior to reversal choices (Fig. 5B), and it exclusively contributed to the preparatory subspace for evoked reversals (Fig.6E, 7D). Consistent with these observations, and contrary to RIB, chemogenetic inhibition of RIS neurons reduced the bias towards reversal response outcomes (Fig. 8C). Conversely, activating RIS acutely via optogenetics for 5 s before the 21% O_2_ stimulus (Fig. S10G) led to more reversal response decisions (Fig. 8F, Fig. S10I).

In summary, not all neurons carrying information about choice bias could be causally linked to final action selection. This finding implies that control over choice bias is a function dedicated to specific neuronal cell classes, while information about choice bias is broadly shared across the nervous system.

## Discussion

Using oxygen chemosensory behaviors in *C. elegans*, we investigated the neuronal circuit dynamics underlying behavioral variability. We developed a binary choice paradigm in which animals were exposed to repeated high-oxygen (21% O₂) stimuli and responded with one of two distinct actions: an immediate pause followed by reversal, or sustained slowing followed by accelerated forward crawling (Fig. 1F). Unlike typical perceptual decision-making tasks, where choice is a function of sensory discriminability ^11^ our assay presents an unambiguous stimulus, thereby isolating behavioral variability arising from intrinsic sources, similar to the variability observed in escape paradigms ^79,80^. Choice bias was not an idiosyncratic property of individual animals but instead varied from trial to trial, with a persistent bias extending over a timescale of minutes or approximately two consecutive trials. Our assay thus serves as a paradigm for variability in instantaneous decision-making, as opposed to variability associated with longer-timescale factors such as developmental history or other lifelong individuality traits. With this framework, we systematically investigated the behavioral and neuronal bases of variable decision-making.

Choices correlated with a preparatory behavioral state: slower locomotion at stimulus onset was associated with reversal outcomes. A similar preparatory slowing behavior can be observed prior to spontaneous reversals ^4,40^. However, we found no evidence for a causal relationship between locomotion speed and choice, as manipulating speed at the level of motor neurons for forward crawling did not alter choice bias. This finding indicates that choice variability is not a consequence of biomechanical constraints or initial behavioral conditions propagated through a bottom-up signal; rather, it arises via a top-down mechanism originating from neuronal circuits upstream of motor execution, which likely shape both the preparatory behavioral state and the subsequent choice in parallel. In this respect, our paradigm is distinct from food-lawn leaving decisions in *C. elegans* that can be predicted and controlled via behavioral state: they are correlated with a high-speed behavioral roaming state and can be induced by suppressing feeding behavior directly via the pharyngeal feeding muscles ^54^.

To describe the sensory-to-motor transformation process at the neuronal level, and to subsequently identify the circuit mechanisms governing choice, we employed whole-brain calcium imaging. This approach faithfully captured the full transformation process, including sensory neuron activation, fictive preparatory slowing, and reversal commands, recapitulating the trial-to-trial choice variability observed in freely behaving animals (Fig. 4A–E). Notably, these experiments were performed in immobilized animals, demonstrating that the qualitative steps of sensory-to-motor transformation— including choice variability—do not require feedback from successful behavioral execution.

Brain-wide activity revealed a layered sensory-motor architecture with auxiliary neuromodulatory neurons: two sensory layers comprising primary sensory neurons and a small set of interneurons reliably encoded the external stimulus. Notably, these sensory interneurons (PVP, AUA, RMG) are known synaptic partners of primary O_2_ sensory neurons. The sensory responses in neuromodulatory neurons ALA, RID, and RIS were unexpected and presumably occur via indirect synaptic communication or extrasynaptically. Moreover, two motor layers, comprising premotor interneurons and motor neurons, were primarily locked to a brain-wide motor command cycle ^4^. At this layer, the main effect of the stimulus was a reliable downmodulation of forward interneurons like RIB, as well elevating the transition probability from the forward motor command to the backward motor command. These results reveal a clear segregation between a sensory domain, encoding the environment, and a motor domain encoding the current behavioral state. These results are consistent with connectome-based analyses suggesting a modular, layered architecture of the *C. elegans* nervous system, in which sensory layers segregate from premotor and motor layers ^72,73^.

Both domains, nevertheless, interact with each other. Comparing stimulus-evoked and spontaneous motor command states revealed that sensory circuits recruit qualitatively the same neuronal population dynamics as spontaneously initiated actions; however, we observed stimulus-dependent amplitude modulations in individual neuron classes, likely reflecting stimulus-specific tuning of behavioral parameters such as locomotion speed (Fig. 4F-J). Crucially, however, our findings demonstrate that sensory-to-motor transformation is not a simple feed-forward reflex pathway but occurs at the intersection of intrinsic brain dynamics and externally evoked activity patterns. While sensory circuits reliably perceive and encode external stimuli, sensory-evoked behaviors are generated within the context of brain-wide neuronal population dynamics that also correspond to spontaneous, uninstructed behaviors. These motor dynamics are largely pre-patterned within the nervous system and are recruited by sensory inputs to produce evoked behavioral responses, suggesting that the role of sensory processing is not to construct motor commands in a bottom-up fashion, but to engage and shape pre-existing motor repertoires. In conclusion, our data reveal a processing hierarchy in which sensory information is reliably encoded by dedicated sensory circuits; however, further downstream, motor commands are distributed across neurons that multiplex sensory and motor signals exhibiting both behavior-modulated and sensory-modulated activity. This architecture suggests that behavioral dynamics are an intrinsic property of the nervous system, with action sequences that are pre-patterned and can be initiated spontaneously yet are readily recruited by sensory circuits to generate stimulus-driven responses.

To identify neuronal correlates predictive of choice prior to stimulus onset, we employed tailored machine-learning approaches. Sensory neurons exhibited stereotypic response patterns devoid of information regarding choice bias, precluding the possibility that choice variability arises from external noise in sensory stimuli or from variability in sensory perception. We therefore conclude that sensory circuits reliably report environmental conditions, while behavioral transformations—including motor command generation and choice bias must occur further downstream. Analogous findings have been made in olfactory neurons reliably responding to odorant stimuli while triggering a variable response to terminate reversals ^81^. Notably, these observations were enabled by the use of strong ex-afferent stimuli operating in open loop with the animal’s current behavioral state. In other settings, for example, when animals navigate chemosensory gradients, sensory perception operates in closed loop with ongoing behavior, such that variations in perception become coupled to underlying variability in locomotion. Future studies should investigate how sensory decision-making operates under such closed-loop behavioral conditions.

Unlike sensory neurons, individual interneuron classes exhibited activity patterns that segregated between the two future choice outcomes during the forward state prior to stimulus onset (Fig. 5B, Fig. S6). Among these were the sensory interneurons AUA and PVP, the proprioceptor DVA, the arousal-related neuromodulatory neurons RIS and RID, and the premotor interneuron RIB. Single-neuron linear discriminant analyses showed that these differential activity patterns carried some predictive power regarding future choice outcome, suggesting their involvement in the decision-making process. In general, our single-neuron models performed comparably to the sole successful single-behavior model, which was based on locomotion speed. The top-performing neuronal model used RIB activity; notably, RIB neurons have been previously implicated in the regulation of forward locomotion speed ^4,48^. Our results suggest that RIB serves a dual function: regulating crawling speed to establish a preparatory slowing state while simultaneously biasing the neuronal network toward initiating a future reversal response. Among the top-performing neurons was also RIS, which is a sleep-active neuron and involved in sleep regulation ^66,78^. During wakefulness, RIS activity can transiently increase during preparatory slowing prior to reversal events ^45^, in contrast to RIB, whose activity declines in these instances ^4^. Our previous work demonstrated that atmospheric O₂ serves as an alarming wake-up signal for sleeping *C. elegans* ^66^. Taking these observations together with the characteristic RIS activity signature in brain states predictive of upcoming reversals, we speculate that this may reflect a state of low arousal in which RIS functions to maintain or even elevate responsiveness to O₂ stimuli.

Moreover, a joint neuronal population model outperformed any single-neuron-based decoder, showing that information about choice bias is widely distributed across the nervous system. Critically, our population-based choice model revealed slowly evolving neuronal state trajectories that diverged between outcome categories tens of seconds before stimulus onset. These data indicate that the initial conditions influencing variable behavioral decisions reflect deterministic, slowly evolving neuromodulatory states rather than fast stochastic fluctuations in neuronal activity or synaptic transmission ^29^. In these respects, our experimental paradigm differs further from the variable olfactory response paradigm reported by Gordus et al., where reversal termination was counteracted by the same neurons implicated in generating the reversal motor command, rendering the current behavioral state more resilient to the sensory input ^81^. Unlike in the present study, we describe a prior neuronal state that sets initial conditions for future choices.

We next generated another PLS model that enabled us to predict future spontaneous reversal events out of the forward locomotion state. The corresponding neuronal subspace overlapped with the one predicting evoked reversals (Fig. 7C; compare Fig. 6E with S9A). Both states receive the strongest contribution from interneurons and motor neurons active during forward locomotion. These neurons typically decrease their activity prior to reversal onset and thereby mediate the preparatory slowing state ^4,47^. However, the subspace predicting evoked reversals appears shifted towards stronger modulation in a subset of interneurons like RIB, DVA, and the yet largely uncharacterized class PVN, and received unique contribution from the neuromodulator neuron class RIS (Fig. 7E). Thus, our analyses revealed quantitative and qualitative distinctions between both subspaces. Choice bias can only be partly explained by a preparatory behavioral or neuronal state in which the nervous system is already primed to transition spontaneously between actions. Moreover, choice bias is additionally influenced by distinct neuronal classes, particularly those involved in neuromodulation and arousal. These observations are conceptually important, as they suggest that neuronal subspaces encoding related task parameters must not be discretely separable but can be flexibly modulated. Crucially, RIS activity was rising slowly prior to the onsets of 21% O_2_ stimuli. The amplitude of this rise contained information of future choice outcome. This indicates that one source of variability in this system is variation in the activity dynamics of RIS, which is then further correlated with a pronounced motor preparatory state for forward-backward transitions. While RIS activity was entrained by the stimulus pattern in this study, it exhibits also spontaneous activity dynamics during forward states ^66,45^, suggesting that RIS mediated biasing of choice outcome must not be solely sensory derived. Interestingly, RIB is a major postsynaptic partner of RIS^33,36,37^, suggesting a channel by which an initial variation could propagate through the network.

Finally, guided by our statistical analyses, we selected individual neuronal classes to test for causal involvement in choice bias. We found that either inhibition or excitation of the neuromodulatory neuron RIS or the interneuron RIB influenced choice bias, whereas the same manipulations in B-type motor neurons or DVA interneurons did not. Notably, all these neuronal classes contributed to the population model predicting future choice. These results reveal a dissociation between representational features of global brain activity, by which information about choice is widely shared in a distributed manner, and causal circuit elements, through which choice bias can be controlled only via a specific subset of neurons within that distributed population. We nevertheless propose that the distributed sharing of cognitive variables such as choice bias across the brain is functionally meaningful. In the case of B-type motor neurons, we suggest that choice-related modulation underlies the preparatory slowing state, which likely facilitates a more efficient biomechanical transition from forward to backward crawling. Distributed modulation of other neuronal classes by choice bias may serve additional diverse functions; for instance, providing behavioral context for sensory integration of complex stimuli in natural environments, which could be investigated in future studies.

Typically, studies of decision-making in mammals introduce learned delay periods to temporally segregate the circuit dynamics underlying decision formation, motor preparation, and motor execution ^11^. However, in most natural settings, animals cannot rely on such a fixed task structure but must make rapid decisions on an ongoing basis, requiring the integration of cognitive processes with the neuronal dynamics governing ongoing behavior. Here, we developed a combined analytical and experimental framework to disentangle these processes in the context of an instantaneous behavioral decision. Leveraging the tractability of an invertebrate nervous system, we show that decision-making, motor preparation, and motor execution recruit overlapping circuit dynamics and are therefore deeply intertwined at the neuronal level; yet all three processes can be analytically distinguished and causally dissociated through targeted manipulations of specific neuronal classes.

The functional localization and specification of brain regions, neuronal circuits, and cell types is a foundational neurobiological principle established across invertebrates, vertebrates, and mammals, including humans ^17,82^. This is further corroborated by optogenetics, where activation of individual circuit elements can trigger complex behaviors or cognitive states ^83,84^. Yet recent discoveries of distributed information encoding across the brain present a fundamental challenge to this classical view ^85^. Here, based on our findings in a tractable *C. elegans* model, we propose a generalizable framework that reconciles these two seemingly contradictory concepts. We posit that while individual circuit elements execute their specialized functions locally, information about the corresponding behavioral and cognitive variables is simultaneously broadcast across the brain, providing essential context for other local neuronal computations. This architecture i.e., specialized functions embedded within a globally informed network, may represent a universal principle of neuronal computations.

## Acknowledgements

The computational results of this work have been achieved using the Life Science Compute Cluster (LiSC) of the University of Vienna. The authors would like to thank Liana Akobian for establishing a data pre-processing and imputation pipeline, Charissa Murphy for experimental help, Julia Riedl and Pietro Verzelli for critical reading of the manuscript. The research leading to these results has received funding in part by the Austrian Science Fund (FWF) 10.55776/COE16, by the Vienna Science and Technology Fund (WWTF) [Grant ID: 10.47379/LS23070], from the European Research Council (ERC) under the European Union’s Horizon 2020 research and innovation programme (*elegans*BrainBodyEnvi, #101054527), from the Simons Foundation (\#543069, NC-GB-CULM-00003196-01), the International Research Scholar Program by the Wellcome Trust and Howard Hughes Medical Institute (\#208565/A/17/Z), the University of Vienna, and the Research Institute of Molecular Pathology (IMP).

## Author contributions

J.M., A.P. and M.Z. conceived the study and designed experiments. J.M., C.F. and M.Z. conceived the computational methods. J.M. implemented the computational methods and analyzed the data. C.F. guided the data analyses. M.Z. led the studies. J.M., C.F. and M.Z wrote the manuscript.

## Methods

### Worms

For all experiments, young adult worms (with 4-12 eggs) cultured at 20^0^C on food plates (Nematode Growth Media (NGM) plates seeded with *Escherichia coli* OP50) were used. Worms with the *lite-1 (ce314)* mutation for reduced light responses ^86^ are used in this study. For all strains and genotype details, see supplementary table S1.

### Preparation of agarose surfaces for behavioral assays

The behavioral assay protocol is primarily designed by keeping in mind that it should closely mimicking our immobilized neuronal imaging conditions. First, to compensate for food deprivation due to muscle paralysis in the imaging worms, worms were food-deprived for 30 minutes before the behavior recording (optimized feeding state) by picking them to a new NGM-agar plate without food. Second, the immobilized worms inside a microfluidic device have restricted food availability due to the absence of pharyngeal pumping. So, we limited the availability of food to the worms in the behavioral assay by using an extremely diluted food patch (optimized food availability). This was achieved by starting from a food-buffer solution (OP50 in NGM buffer) of 1.6 OD and then diluting it to a working concentration of 1:100. This food solution was prepared freshly on each recording day. We used 10 cm (diameter) NGM-agar (1.75%) plates with a 30 mm x 30 mm square arena in the middle, confined with Whatman filter paper as the behavioral assay plates. The Inside of the arena was uniformly seeded with 200 µL of the diluted food solution for roughly 30 minutes to make sure the liquid was soaked into the agar before the recording started. All the inside borders of the Whatman filter paper were soaked with 20 mM CuCl_2_ as a repellent to prevent the worms from escaping the arena. For all behavioral assays (including all the chemogenetic and optogenetic experiments), we followed this standardized protocol for consistency.

### Reconstructed gas stimulus protocol

To be consistent with the neuronal imaging condition, for behavioral assays, where O_2_ stimuli are delivered via gas flow, we used a reconstructed gas stimulus protocol that matches the relatively slow diffusion profile through the PDMS layer of the microfluidic device in our imaging apparatus ^41,64^. For this, we first measured the oxygen diffusion using an oxygen-quenching dye, Tris (4,7-diphenyl-1,10-phenanthroline) ruthenium (II) dichloride complex ^87^ (Absorbance: 455nm, Emission: 610-630nm). The dye was solubilized in ethanol and used at a concentration of 10µM. It was loaded into the worm chamber of the microfluidic device and, using an epifluorescence microscope, we measured the fluorescence intensity change at 488nm while varying oxygen concentration (as described for neuronal imaging below), using Fiji software ^88^. We created a calibration curve by measuring the fluorescence intensity across different oxygen concentrations and performing an exponential fit. Then, this calibration curve was used to derive oxygen concentration with our 21-11 stimulus profile. A denoising smoothing using Total Variation (TV) Regularization was applied to reconstruct the stimulus profile. Thus, this simulated oxygen profile (Fig. S1A) was used for all behavior recordings (including all the chemogenetic and optogenetic experiments).

### High-resolution behavior recordings

The worm was placed in the center of the arena on a behavioral assay plate, and a constant 11% O_2_ (mixed with 89% N_2_) was delivered at a flow rate of 50 mL/min. A static lab supply of O_2_, N_2_, and CO_2_ was controlled using a red-y smart GSC -A9TA-BB22 mass flow controller (Vögtling Instruments) for each gas type. These were combined, and the mixed gas was delivered through a transparent plexiglas device with a flow arena of 39 mm x 39 mm x 0.7 mm placed on top of the assay arena, as described previously ^4,76,89^. The stage was illuminated with an infrared LED, and an ROI of 1500 pixels x 1500 pixels (∼1 pixel = 1 µm) was recorded at 10 fps for a total duration of 31.5 minutes, using a custom rebuild of a high-resolution behavior recording system ^90^. Briefly, the 3.125x magnification of telecentric optics was achieved by combining a Raynox MSN-202 lens as the objective with a Raynox DCR-250 Lens as the tube lens. The system was set up as a transmission-light microscope with a 730nm Thorlabs M730L5 LED as back illumination and a Thorlabs ED1-S50 diffuser as a condenser. For imaging, an IDS uEye UI-388xCP-M camera was used. The microscope was controlled by Micro-Manager ^91^ with custom plug-ins for tracking, position control, and time-synchronized gas supply. The tracking was performed by calculating the moments of the image and controlling the x-y motor speed to keep the weighted center of the image in the middle of the camera ROI. All recordings were later converted to video files (AVI files) using custom Python code.

### Histamine inhibition

To inhibit neurons transiently, we used a chemogenetic approach by extra-chromosomally expressing Histamine-gated Chloride channels in a cell-specific manner ^62^. We followed our behavioral assay protocol (see above) but supplemented 40 mM histamine (histamine dihydrochloride, Sigma-Aldrich) to the food-NGM buffer and the NGM plates used for both starvation and recordings. For the His (-) control, instead of histamine, an equal amount of solvent (water) was used.

### Optogenetic experiments

To acutely activate specific neurons, we applied an optogenetic approach with worms expressing a red-shifted, codon-optimized channelrhodopsin, Chrimson ^92^, in a cell-specific manner. 20-24 hours before the experiment, L4-stage worms were picked onto fresh NGM plates seeded with OP50 plus either 200 µM all-trans retinal (ATR, from Sigma-Aldrich R2500) or an equal volume of the solvent (ethanol) lacking ATR (Control). We followed our optimized behavior assay protocol mentioned above, but the worms were starved and recorded in the dark. We used 591nm light, at ∼0.4mW/mm^2^ for optogenetic stimulation. In our experimental design, the same gas stimuli were used (16 trials of 11%-21% O_2_ stimuli) but now coupled to an opto-stimulus. i.e., optogenetic activation preceded each of the 21% O_2_ stimuli (see Fig. 8D-F, Fig. S10D, G). For RIB and DVA neurons, this was a pulsed light (20Hz) stimulus for 15 seconds, and for the RIS neuron, it was a constant light for 5 seconds. These timings were empirically determined in test recordings aimed at visible, reliable, and sustained behavioral effects in each transgenic line. Our optogenetic recording setup differs from the high-resolution behavior setup explained before in the following. An optogenetic stimulation was incorporated using a dichroic mirror through the objective. For precise tracking of animal position, we estimated the nose position using DeepLabCut (DLC) ^93^ based on live inference (nose tracking) using Python-based custom code. We recorded an ROI of 1104 pixels x 1104 pixels (∼1 pixel=1.82 µm) at 50 fps, using a high-speed camera (25GigE Area-Scan Cameras BOLT Series, HB-8000-SB-M, from Emergent Vision Technologies). Along with the recording settings, the gas stimulus protocol was also executed using custom-written Python software.

### Immobilized whole-brain Calcium imaging

For immobilizing worms while imaging, we used a non-invasive method of muscle inhibition. We generated worms with Histamine-gated Chloride channels expressed pan-muscularly with the *Pmlc-2* promoter ^62,65^. Additionally, they expressed a pan-neuronal, nuclear-localized GCaMP6f ^63,94^. These worms were incubated on an OP50-seeded NGM agar plate mixed with 20 mM histamine for 30 minutes before imaging. For the imaging, we used two-layer PDMS microfluidic devices designed and manufactured as described previously ^4,63,64,95^. This microfluidic device was connected, and the worms were loaded as previously explained ^63^. The buffer for loading and imaging in the microfluidic device was a food solution (*E. coli* OP50, diluted 1:100 in NGM buffer from a 1.6 OD stock), containing 20 mM histamine.

We acquired high-resolution whole-brain Ca^2+^ recordings as previously described ^4,63,66,67^. Shortly, we used an inverted spinning disk confocal microscope (Zeiss Axio Observer.Z1 with attached Yokogawa CSU-X1) with an EMCCD camera (Andor iXon DU897_BV 3972) and a 40x 1.2 LD LCI Plan-Apochromat water-immersion objective (Zeiss). We performed volumetric image acquisition with a 20 ms exposure time and 2μm steps between Z-planes, achieved using a Piezo stage (P-736 PInano, Physik Instrumente GmbH). These settings allowed us to image approximately 14-15 planes at 3 volumes per second for a total duration of 31.5 minutes. We used a blue laser (LBX-488nm-100-CSB) at a power of 2.3-4 mW/mm^2^ for GCaMP excitation. The gas stimulus protocol was given using a gas mixer with mass flow controllers (Vögtling Instruments), which was controlled along with the imaging settings via custom-written Micro-Manager software. A constant gas flow of 11% O_2_ (mixed with 89% N_2_) at a 50 mL/min flow rate was maintained while setting up each imaging experiment.

## Data Analysis

### Behavior analysis pipeline for behavioral annotation

We used a custom analysis pipeline to extract the head-to-tail centerline and full-body curvature (kymograms) from our high-resolution behavioral recordings ^47^. These outputs were subsequently used to annotate different behavioral states, including forward locomotion, reversals, and dorsal and ventral turns. Briefly, the raw images were first binarized to generate a worm mask using a U-Net-based neural network, a deep learning architecture widely used for image segmentation tasks ^96^. To accurately reconstruct challenging body postures, such as coiled worms, we independently trained a second U-Net specialized for these complex morphologies. Next, we used DeepLabCut, an open-source toolbox for markerless animal pose estimation and tracking, to identify the nose and tail positions of the worm ^93^. These landmarks served as anchor points for fitting a centerline consisting of 100 uniformly spaced spline segments through the center of the binary mask using the SciPy library in Python ^97^ (Fig. 1B). The dorsal and ventral sides of each animal were manually assigned based on the position of the vulva. Using the resulting centerline coordinates, body curvature was calculated for each segment as previously described ^47^, yielding curvature kymograms that describe body posture over time (Fig. 1B). Eigenworms were subsequently computed by performing principal component analysis (PCA) on the body posture data ^98^. To minimize distortions caused by rapid head movements and the often incompletely segmented transparent tail, which typically produces noisy centerlines at both ends of the animal, the first and last 20 body segments were excluded from the PCA. Forward and reversal states were then automatically identified from the phase relationship between the first and second eigenworm projection amplitudes ^47,98^. These automated annotations were subsequently manually curated to remove false reversal detections and to correct for missing or erroneous annotations arising from unresolved coiled body postures.

Turning events were detected as previously described ^47^. Based on the manually annotated vulva position (left or right side of the animal), curvature values were assigned a sign, with ventral curvature defined as positive and dorsal curvature as negative. This signed curvature was used to detect turning events and classify them as either dorsal or ventral. A turn was defined as beginning immediately after the end of a reversal and continuing until the summed head curvature (body segments 1–10) changed sign. Head amplitude was defined as the maximum curvature of the head region, calculated as the peak curvature across body segments 1–10.

Slowing states were identified from changes in forward locomotion speed. First, forward speed was smoothed using a 3-second moving average. The temporal difference of this smoothed speed trace was then calculated, and a transition to negative values was defined as the onset of a slowing episode. To reduce false-positive detections caused by small fluctuations or tracking jitter, only continuous periods of decreasing forward speed lasting at least 1 second were classified as slowing states.

### Behavioral response classification

Each trial was manually assigned to one of three response categories according to the animal’s behavior within predefined time windows relative to stimulus onset.

#### Reversal response

A reversal was initiated within the response window (0–8.5 s after stimulus onset).

#### Slowing response

No reversal was detected within the response window, but the animal exhibited a decrease in forward locomotion speed during this period.

#### Already reversing

The animal had already initiated a reversal during the 2 s immediately preceding stimulus onset (−2 to 0 s) and was therefore excluded from response classification.

#### Missing data

Due to tracking failure.

### Behavioral features extraction

#### Speed

The worm’s speed was calculated from the centroid trajectory. For high-resolution behavior recordings, centroid positions were obtained from the real-time stage position recorded during camera tracking. For nose-tracking behavioral recordings (optogenetic experiments), centroid coordinates of the binary worm mask in each frame were calculated with respect to the real-time stage position (nose position). Speed outliers were removed by clipping values exceeding 0.3 mm/s, and the resulting speed trace was smoothed using a 10-frame (1 s) moving average. During analyses of forward locomotion speed, time points corresponding to reversal states were excluded.

#### Angular speed

Angular speed was calculated as the complement of the angle defined by three consecutive points along the worm’s crawling trajectory ^56,57^. The crawling trajectory was obtained from the centroid positions after smoothing with a 3-frame moving average. Angular speed therefore provides a measure of the rate of change in the worm’s direction of movement.

#### Curvature

Unless otherwise stated, body curvature refers to the total body curvature, calculated as the sum of the absolute curvature values across all 100 body segments. Curvature at each segment was defined as the reciprocal of the radius (*R*) of the osculating circle fitted to the centerline at that segment:

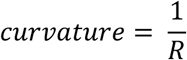

Forward run length: Forward run length was defined as the elapsed time spent in the forward locomotion state since the most recent reversal event.

Head bending frequency: The worm’s centerline was represented by 100 uniformly spaced segments (Fig. 1B). Head curvature was calculated as the sum of the absolute curvature values of the first 10 body segments. Each transition in the sign of the head curvature (positive to negative or vice versa) was counted as one head oscillation. Head frequency was then calculated as the number of head oscillations per second. A continuous time series is generated from this by taking a rolling mean over 10 frames (1 s).

Roaming/dwelling state: Previous studies have shown that foraging *C. elegans* alternate between two long-lasting locomotory states: roaming and dwelling ^55–58^. The dwelling state is characterized by local search behavior with low crawling speed and high angular speed, whereas the roaming state corresponds to global exploration with high crawling speed and low angular speed ^55,56,63^. Following the approach described previously ^56,57^, 10-second binned values of speed and angular speed were classified into roaming and dwelling states using the decision boundary (y = x/150). This classification yielded a binary behavioral variable indicating the locomotory state of the worm (roaming = 1; dwelling = 0), which was used as a behavioral feature.

### Shuffle trigger control for the empirical cumulative distribution functions (ECDFs)

To verify that the stimulus-triggered empirical cumulative distribution functions (ECDFs) reflected genuine stimulus-dependent dynamics rather than the temporal structure of the dataset, we generated a shuffled-trigger control. For each animal, 16 trigger points were randomly selected within the 11% O₂ period, matching the number of experimental stimulus presentations. For each randomly selected trigger, a 30-second analysis window was extracted, and the ECDF was calculated using the same procedure as for the stimulus-triggered data. This randomization procedure was repeated 1,000 times, and the mean shuffled ECDF was used as the control.

### Neuronal traces/time series extraction

Neuronal time series were extracted as described in detail in Kato *et al*., 2015, using custom MATLAB scripts ^4^. Single-cell fluorescence intensities (F) were computed by tracking the intensity maxima in each volume over time after subtracting the background. ΔF/ F_0_ was calculated for each neuron using the mean fluorescence intensity across time as F_0_.

### Neuron Identification

In our whole-brain calcium recordings, we roughly detected 105–185 neurons per recording. The identification of the neuronal cell classes (IDing) in these imaging data was done using the logic explained in our previous studies ^4,63,66,67,99^. Briefly, in our datasets, we identified neurons by considering the following: their activity (and correlation to the neurons in the same class), relation to neighboring neurons, shape, and relative position (http://www.wormatlas.org). This way, we only identified the identity of active neurons in the datasets (82 neurons in total, Fig. 3A, Fig. S4A, B). Later, we confirmed the identities of many neurons by separately performing the same experiments using NeuroPAL worms (N=6) ^100^. This added confidence to the neuron IDs. There were still neurons with consistent activity in our datasets, but they did not appear in the NeuroPAL worms. These are highlighted in Fig. S4B.

### 4-phase state annotation and rise onset detection

For each neuronal trace, by thresholding the derivative, we detected peaks using custom-written MATLAB code as described previously ^4^. Based on the detected peaks, traces were color-coded with 4 state phases: low, rise, high, and fall states. A transition between the low and rise state phase was defined as the peak onset and used for response probability and delay time calculations (Fig. 3E-F). Note that, since it is based on peak detection, we limited these analyses to neurons with discrete state transitions. Neurons with ramping activity and thus slow phase changes are not used. Based on several previous studies ^61^ we used the AVA neuron peaks for reversal command state annotation in the immobilized neuronal recordings, as the reversal onsets reliably correspond to the AVA peak onsets in freely moving worms ^4,69,70^, which is used for all reversal response-triggered analyses.

Using the AVA state annotation, we defined the following fictive discrete behavior states (should last at least 1 s) in our immobilized neuronal recordings:

#### Reversal: AVA rise state

Dorsal/Ventral Turns: AVA fall state is defined as a turn state. As shown in previous studies, Dorsal/Ventral identity is determined using the activity of SMDD and SMDV neurons ^4,63^. By comparing the peak amplitudes, we define higher SMDD activity as a Dorsal turn, and higher SMDV activity as a Ventral turn.

Forward: AVA low state.

### Modulation index calculation

We identified two types of neuronal activities in the whole-brain calcium recordings. First, the sensory activity reports the input sensory information of 21% O_2_. Second, there is the motor-related activity, reflecting the ongoing motor dynamics of the worm. Thus, for every neuron, we looked at the sensory stimulus-triggered (change to 21% O_2_) and the AVA rise onset-triggered (reversal-triggered) average traces. We considered only the spontaneous reversals in the stimulus period, obtained from all 11% O_2_ downshift periods (excluding the baseline). Then, in each case, we calculated the min-max difference in a 30-second window (post-stimulus) of the smoothed triggered averages (2-second moving mean, as shown in Fig. 3C). We called these absolute values “stimulus-triggered activity (Stim)” and “reversal-triggered activity (Rev)” of each neuron.

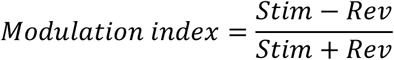

Fig. 3D shows the modulation indices of some active neurons overlaid on their connectome (Full details in supplementary table S2).

### Pre-processing for PLS and PCA model analysis

Neuronal traces used for PLS and PCA models are pre-processed using the following steps (relevant for Fig. 4, 6 & 7):

1. Data filtering: We only retained neurons captured in more than 50% of the recordings to be used as input for all models. In other words, each neuron should have less than 50% missing data. This way, we had 62 neuronal traces as inputs.
2. Scaling: While concatenating neuronal traces from multiple datasets (recordings/animals), we scaled twice to ensure uniformity. Once within each dataset, i.e., each dataset separately. Then, we scaled across datasets, i.e., after concatenating all datasets. We used the RobustScaler function from the sklearn library of Python. RobustScaler (with_centering=False, with_scaling=True, quantile_range= (5, 95)).
3. Handling missing data: We have missing data due to failure in identifying a neuron in some datasets (recordings/animals). These missing data were handled by imputation. We used a probabilistic principal component analysis (PPCA) model (https://github.com/allentran/pca-magic) fitted on the entire dataset for imputing missing neuronal traces ^101^.

### Partial Least Squares (PLS) data preparation and analysis

Partial Least Squares regression (PLS-R) is a multivariate supervised machine learning approach that identifies linear combinations of predictor variables (latent components) that maximally covary with the response variable by projecting both datasets into a shared orthogonal latent space ^102^. PLS-R has been widely applied in neuroscience to relate high-dimensional neural or behavioral data to experimental variables ^102–104^. As a linear regression model, it can be represented as:

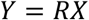

*X* (n, t) = predictor variables, independent variables
*Y* (n, m) = response variables, dependent variables
*R* = regression coefficient

PLS-R finds the latent components by iteratively maximizing the covariance between transformed predictor and response variables using Singular Value Decomposition (SVD):

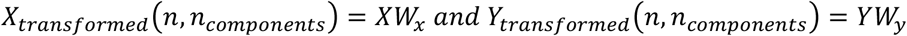

*W_x_* (t, n_components_) and *W_y_* (m, n_components_) are the weight matrices ^102,103^.

Three PLS-R models were developed in this study: (1) the behavioral PLS model, (2) the sensory PLS model, and (3) the spontaneous PLS model. All models were implemented using the PLSRegression function from the Python package scikit-learn ^105^.

#### Behavioral PLS model

For the behavioral PLS model, the predictor variables consisted of the behavioral features extracted from the behavioral recordings (described in the ‘Behavioral feature extraction’ section). For each trial, only the 5-second pre-stimulus period (−5 to 0 s) was used as input. To match the temporal resolution of the calcium imaging data, these behavioral traces were downsampled by a factor of two, resulting in a predictor matrix of (X) (322 trials × 5 s × 5 fps, 6 features). Missing values due to tracking failures were imputed using the mean value across all trials. The response variable (Y) (322 trials × 5 s × 5 fps, 1) was a binary label corresponding to the animal’s response to the sensory stimulus in the post-stimulus response window (reversal or slowing). A PLS-R model with two latent components (n_components=2) was fitted and used for the analyses presented in Figures 2C and 6A.

#### Sensory PLS model

For the sensory PLS model (used in Figures 6 and 7), the predictor variables consisted of the preprocessed neuronal activity traces (62 neurons; see above-mentioned preprocessing steps). Similar to the behavioral PLS model, for each trial, the 5-second pre-stimulus window (−5 to 0 s) was used as input, resulting in a predictor matrix of (X) (117 trials × 5 s × 5 fps, 62 neurons). The response variable (Y) (117 trials × 5 s × 5 fps, 1) was a binary label representing the worm’s behavioral response to the 21% O₂ stimulus (reversal or slowing), determined from the AVA activity state. When directly compared with the spontaneous PLS model, the pre-stimulus window used for input was changed to 8.5 seconds (−8.5 to 0 s) for consistency (relevant for Fig. 7). A PLS-R model with five latent components (n_components=5) was trained to predict the sensory-evoked behavioral response decision and is referred to throughout this paper as the sensory PLS model.

#### Spontaneous PLS model

The spontaneous PLS model (used in Figures 7C and 7D) was developed to predict spontaneous reversals during the absence of sensory stimulation. The predictor variables consisted of the activity of all interneurons and motor neurons (49 neurons), excluding sensory neurons. To maintain consistency with the sensory PLS model, the initial 450-second baseline period under 11% O₂ was excluded, and the same preprocessing pipeline was applied. Spontaneous reversal events were defined by AVA activation during the 11% O₂ downshift periods. Forward events were defined as continuous, non-overlapping 8.5-second forward locomotion epochs occurring during the same downshift periods. These two event types constituted the binary classification labels (spontaneous reversal or forward locomotion). For each event, neuronal activity during the preceding 8.5-second window (−8.5 to 0 s) was used as the predictor matrix, resulting in (X) (1337 events × 8.5 s × 5 fps, 49 neurons). Because spontaneous reversal events occurred approximately eight times less frequently than forward events, the dataset was highly imbalanced. To compensate for this imbalance, a data augmentation strategy was implemented for the spontaneous reversal category. Specifically, the pre-reversal window was shifted by one frame on each iteration to generate eight overlapping input windows per spontaneous reversal event, thereby approximately matching the number of forward events. In effect, the spontaneous reversal samples were augmented eight-fold. The response variable (Y) (1337 events × 8.5 s × 5 fps, 1) consisted of the corresponding binary event labels (spontaneous reversal or forward locomotion). A PLS-R model with five latent components (n_components=5) was trained to predict spontaneous reversal decisions and is referred to throughout this paper as the spontaneous PLS model.

### Linear Discriminant Analysis (LDA)

Linear Discriminant Analysis (LDA) is a widely used supervised machine learning method for classification and dimensionality reduction problems in neuroscience ^106–108^. Briefly, LDA identifies a linear projection that minimizes within-class variance while simultaneously maximizing between-class variance ^59,107^. A simplified explanation of LDA can be illustrated using a one-dimensional, two-class classification problem, where some scalar values must be optimally separated into two classes. The LDA cost function can be represented as the ratio between the between-class variance and within-class variance:

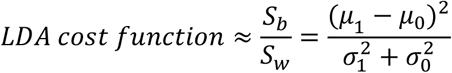

S_b_ = between-class variance, S_w_ = within-class variance
µ = mean of each class
σ = variance of each class

In this study, LDA was primarily used for two applications: 1) classification, 2) dimensionality reduction.

1. For classification, LDA was used to calculate cross-validated prediction accuracy for individual behavioral features (Fig. 2C, 6A) and neuronal features (Fig. 6A, S7), as well as for multivariate PLS models (Fig. 2C, 6A, 7C). LDA was implemented using the LinearDiscriminantAnalysis function from the Python scikit-learn package ^105^. For individual features, a 5-second pre-stimulus window (−5 to 0 s), identical to that used for the corresponding PLS models, was used as input to ensure comparability (Fig. 2C, 6A, S7). For behavioral data, cross-validation was performed using a 90% training and 10% testing split with StratifiedGroupKFold (n_splits = 10, shuffle = True). For neuronal data, an 80% training and 20% testing split was used with StratifiedGroupKFold (n_splits = 5, shuffle = True). Statistical significance of classification performance was assessed using permutation testing, implemented using the permutation test (n_permutations = 1000–2000) function available in the scikit-learn package. To avoid data leakage during cross-validation, preprocessing steps that estimate parameters from the data were either omitted or performed independently within each training fold and subsequently applied to the corresponding test fold.
2. LDA was additionally used for dimensionality reduction by projecting PLS trajectories onto a one-dimensional decision axis. This approach was used to visualize and quantify single-trial neuronal trajectories (described in detail in the Correlation analysis of single-trial neuronal trajectories section; relevant for Figure 6D and Figure S8). The sequential application of PLS followed by LDA is commonly referred to as Partial Least Squares Discriminant Analysis (PLS-DA), a widely used approach in multiple fields including neuroscience and biomedical data analysis ^109–111^.

### Principal Component Analysis (PCA)

PCA is a dimensionality reduction technique widely used to visualize patterns in high-dimensional neuronal data ^112,113^. To compare the spontaneous (at 11% O_2_) and stimulus-evoked dynamics (at 21% O_2_), we used PCA. Similar to the spontaneous PLS model, we excluded sensory neurons from this analysis. This was because sensory neurons were exclusively active (and sometimes covaried with motor neurons and motor-related interneurons) during the stimulus-evoked period, which introduced a trivial difference in dynamics. We followed all the preprocessing steps (see preprocessing steps for models’ session) before fitting the neuronal activities to a PCA model (n_components = 3), which was again an implementation from the sklearn Python package ^105^. (Variance explained: PC1= 28%, PC2=17.6%, PC3=6.1%).

### Correlation analysis of single-trial neuronal trajectories

To investigate trial-to-trial variability in neuronal trajectories within the PLS space, we applied a sequential PLS and LDA analysis approach, which transformed high-dimensional neuronal trajectories into a one-dimensional decision trajectory for each trial.

First, preprocessed neuronal activity data (excluding the baseline period—the first 450 seconds of the 11% O₂ recording—and sensory neurons) from the 5-second pre-stimulus window (−5 to 0 seconds) of each trial were used to fit a PLS model (n_components=5). The resulting latent components from this PLS model, representing neuronal trajectories in the PLS space during the same pre-stimulus window, were then used as input for LDA. LDA projected these PLS trajectories onto a single discriminant axis, generating a one-dimensional Linear Discriminant (LD) trajectory, referred to here as the decision trajectory, for each trial.

To quantify the similarity of decision trajectories within each behavioral response category (reversal or slowing), we calculated the Pearson correlation coefficient between the LD trajectory of an individual trial (restricted to periods when the animal remained in the forward locomotion state) and the average LD trajectory calculated from all trials within the same response category. These individual LD trajectories and their corresponding correlation coefficients are shown in Figure S8, with the summary analysis presented in Figure 6D.

### Weighted feature importance calculation

To estimate the contribution of individual neurons to the latent components (PLS1, PLS2, PLS3, etc.) of the sensory PLS model, we first looked at the neuronal loadings of each component separately. To obtain a single measure of each neuron’s overall contribution to the entire PLS model, we combined the component loadings by weighing them according to the importance of each latent component.

In a PLS model, the latent components are ordered according to the amount of covariance they explain between the predictor variables (X) and the response variable (Y). Consequently, earlier components (e.g., PLS1) contribute more to the model than later components (e.g., PLS2 and PLS3) and should therefore receive a larger weight when estimating overall neuronal contributions.

To determine the importance of each latent component, we treated the PLS components as input features for the downstream LDA classifier and calculated the decrease in cross-validated LDA prediction accuracy by permuting each component independently (100 permutations per component). For this purpose, we used the permutation feature importance method implemented in the Python scikit-learn package ^105^ a widely used machine learning approach for estimating feature importance. The average reduction in prediction accuracy was used as the importance score for that component. Using this approach, the importance of the first three components was estimated as: PLS1 = 18.5±0.3%, PLS2 = 11.1±0.3%, and PLS3 = 4.2±0.2%. These component importance scores were then used as weights of the neuronal loadings to calculate the weighted neuronal contributions shown in Figure 6E. Briefly, the loading of each neuron within a given component was first normalized by the sum of the absolute loadings of all neurons in that component. The normalized loading was then multiplied by the importance score (weight) of the corresponding component to obtain the weighted loading of each neuron.

For example, to calculate the weighted contribution of neuron N to component x: Normalized loading of neuron n to component x:

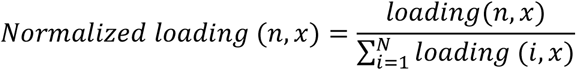

Weighted loading of neuron n to component x:

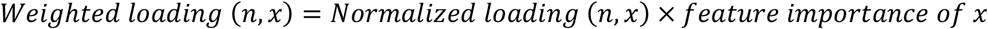

Weighted contribution (%) of neuron n to component x:

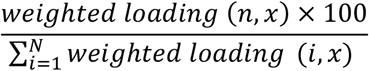

### Neuronal data reconstruction using PLS models

To estimate the extent to which an individual neuron’s ability to predict sensory-evoked reversals can be explained by its ability to predict spontaneous reversals, we performed the following analysis. First, we reconstructed the neuronal activity from the spontaneous PLS model by applying the inverse transformation, which projects the latent variables back into the original high-dimensional neuronal feature space:

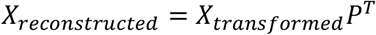

X_reconstructed_ (n, n_features) = reconstructed neuronal traces from the spontaneous PLS model
X_transformed_ (n, n_components) = latent variables of the spontaneous PLS model
P (n_features, n_components) = loadings matrix

This reconstruction represents the component of each neuron’s activity captured by the spontaneous PLS model and therefore acts as a filter for spontaneous activity. We refer to this reconstructed signal as the ‘spontaneous activity’ of each neuron. Similarly, neuronal activity was reconstructed using the sensory PLS model, yielding the ‘sensory activity’ of each neuron, which represents the component of neuronal activity captured by the sensory PLS model.

To compare each neuron’s contribution to spontaneous and sensory-evoked reversal prediction, we analyzed stimulus-triggered averages of these reconstructed neuronal activities. After smoothing the reconstructed traces using a 2-second moving average, stimulus-triggered averages were calculated separately for reversal and slowing trials for both PLS model reconstructions. For each neuron, the maximum difference between the reversal and slowing averages was then calculated within the 20-second pre-stimulus period (−20 to 0 s). This maximum difference was used as a measure of the neuron’s spontaneous and sensory contributions (Fig. 7D). Statistical significance was assessed around the time of the maximum difference using a custom permutation test, and the resulting values are reported in Figure 7E.

### Normalized reversal response probabilities for optogenetic experiments

To quantify how optogenetic stimuli influenced O_2_-evoked reversal response probabilities, we performed control experiments in which optogenetic neuronal activation was applied without the gas stimulus to measure how optogenetic activation alone affects reversal probability in the subsequent time period for the O_2_ stimulus (Light-Only, see Fig. S10D&G). For this purpose, Light-only evoked response probabilities were calculated by taking the mean reversal probability separately for each group (ATR− and ATR+). These means were then subtracted from the respective response probability of each animal in the corresponding experimental condition (Light-Gas) (see S10E, F, I) to calculate the normalized reversal response probabilities used in Fig. 8D-F. All data points falling below zero after normalization were not plotted in Fig. 8D-F but were used for mean calculation and statistics.

### Statistical comparison

Unless specified, we used the Mann-Whitney U test from the SciPy library ^97^ in Python for statistical comparison. For multiple comparison correction, we used the FDR (False Discovery Rate) implementation from the statsmodels library ^114^ in Python.

### Custom permutation test

For statistical analyses involving data pooled across trials, we used a custom permutation test. Response condition labels were randomly shuffled to generate a null distribution of effect sizes (permuted effects), while preserving the number of observations in each condition. This procedure was repeated 2,000–5,000 times, and the observed effect (true effect) was compared with the null distribution to estimate the probability of obtaining an effect of equal or greater magnitude by chance. The resulting permutation *p*-value was calculated as the proportion of permuted effects that were at least as extreme as the observed effect. Results were considered statistically significant at *p* < 0.05 ^67^.

## Declaration of interests

The authors declare no competing interests.

## Declaration of generative AI and AI-assisted technologies in the manuscript preparation process

During the preparation of this work, the author(s) used ChatGPT and Claude for optimizing text expressions and grammar of the manuscript. ChatGPT was used to assist in writing analyses code. The author(s) reviewed and edited the output as needed and take full responsibility for the content of the published article. All thoughts, ideas, and interpretations communicated in this manuscript solely originated from the authors.

---

**Figure S1:**
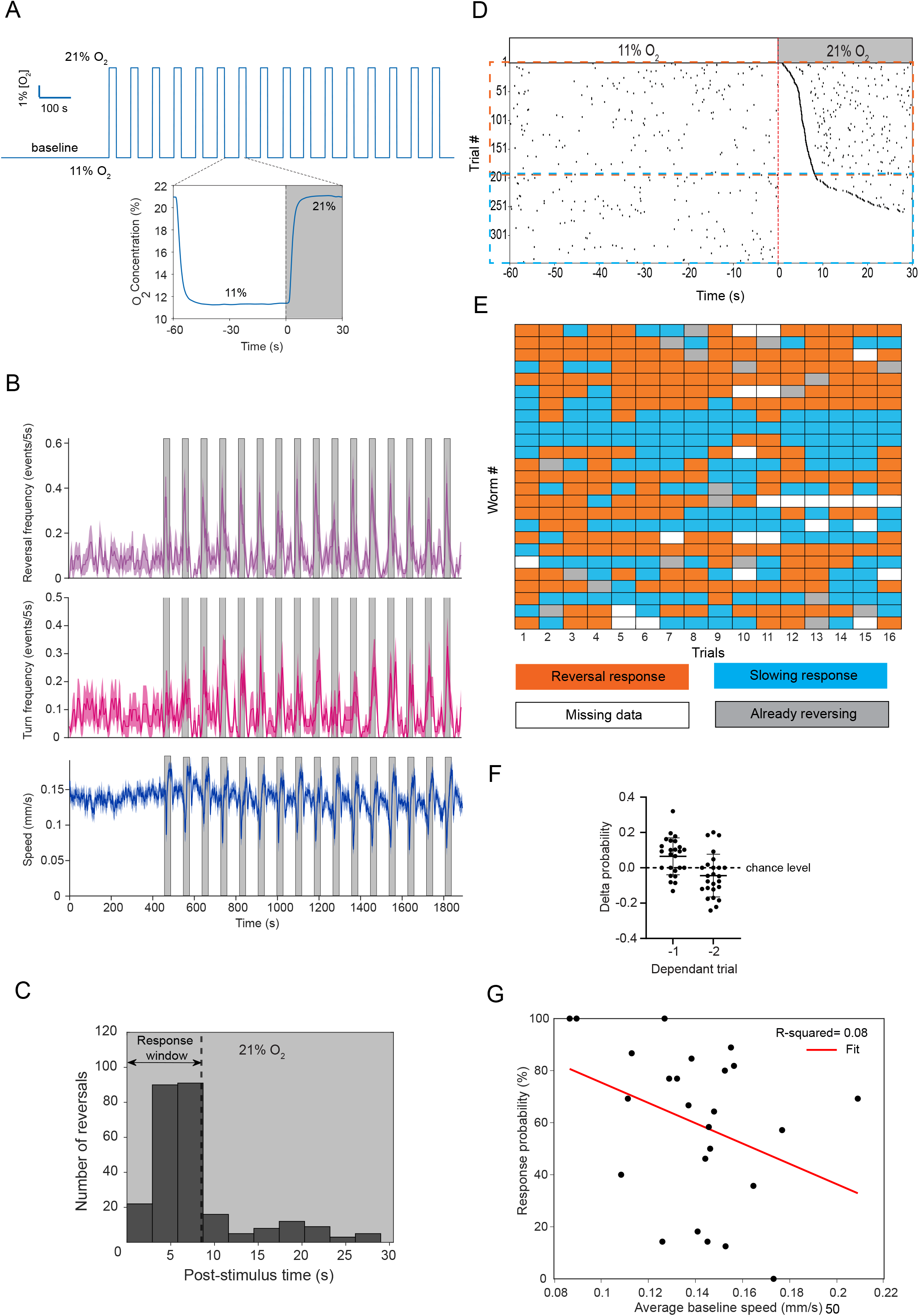
21% O2 stimuli and characterization of behavioral responses of worms, related to. Figures 1 **and 2. A.** Oxygen stimulus profile in a microfluidic (imaging) device, simulated using oxygen quenching dye (see methods). **B.** The average forward speed, reversal probability, and turn probability across the entire recording. (mean±SEM). **C.** Histogram of the first reversal onset time in the 21% O2 (post-stimulus) period. The black dotted line shows the defined response window (0-8.5 seconds). **D.** Stimulus-triggered occurrence of reversal onset events in all trials, sorted based on the reversal response delay within the 21% O2 stimulus period. Orange = reversal response trials and Blue = slowing response trials, N= 25 / 322 (worms/trials). **E.** Response category of each trial. Orange = reversal response, Blue = slowing response, Gray = already reversing, White = missing data due to tracking error. **F.** Probability of the same response in consecutive (-1) and the trial before that (-2). Each dot represents an animal, and the y-axis shows the difference from its shuffle control. **G.** Average baseline (first 450 seconds of 11% O2) speed is plotted against their reversal response probability. Each dot represents an animal. The solid red line shows the best linear fit.

**Figure S2:**
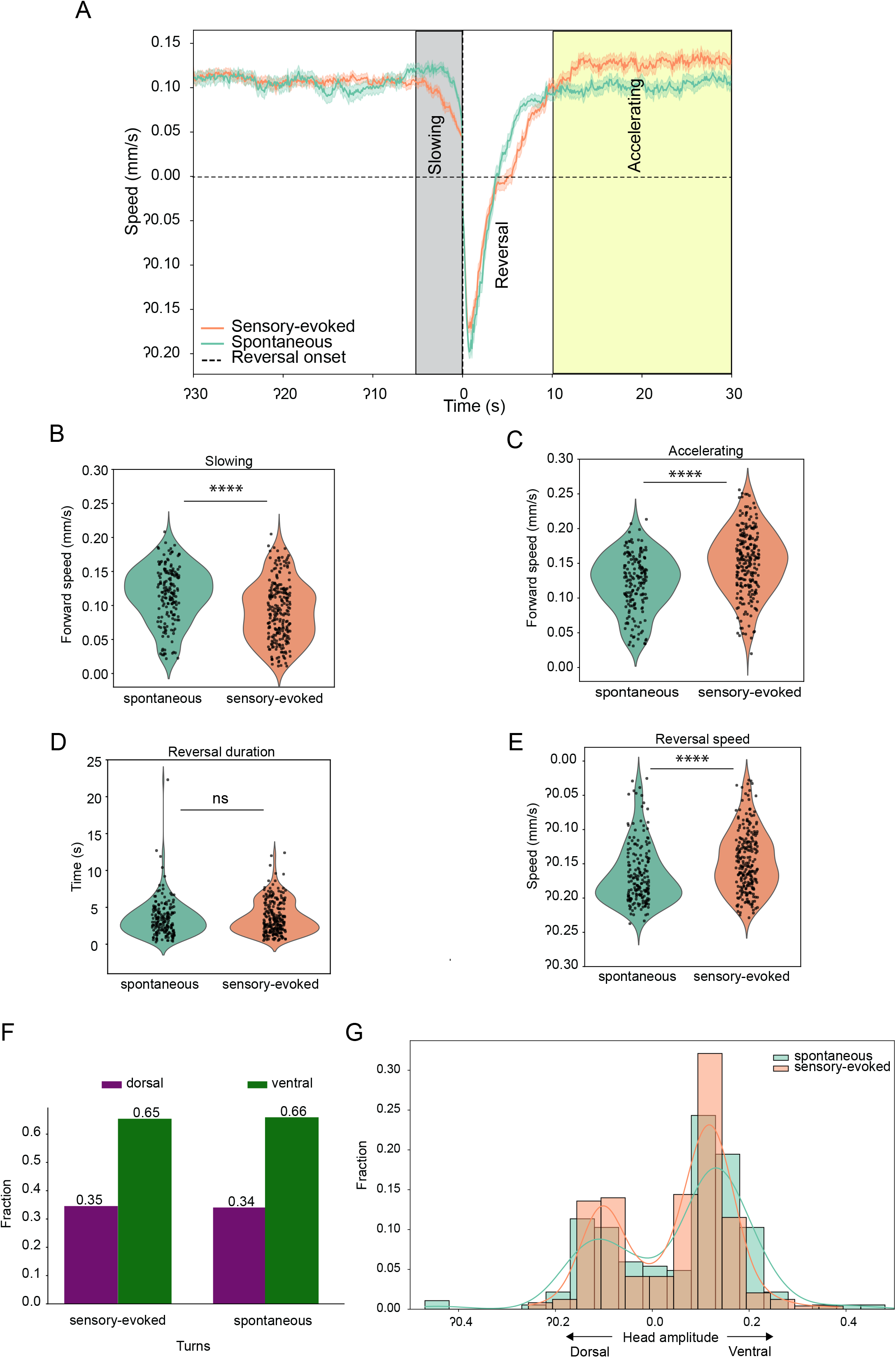
Comparison of spontaneous and evoked behavioral dynamics. **A.** Triggered-average speed in spontaneous (at 11% O2) and evoked (at 21% O2) reversals (mean±SEM). **B-E.** Violin plots of slowing (B), accelerating (C), reversal duration (D), and reversal speed (E). Time windows of slowing (gray) and accelerating (yellow) are as indicated in Fig. S2A. Each dot represents a reversal event. **F.** Fraction of dorsal ventral turns following each reversal category. G. Head swing amplitude during turns after sensory-evoked and spontaneous reversals.

**Figure S3:**
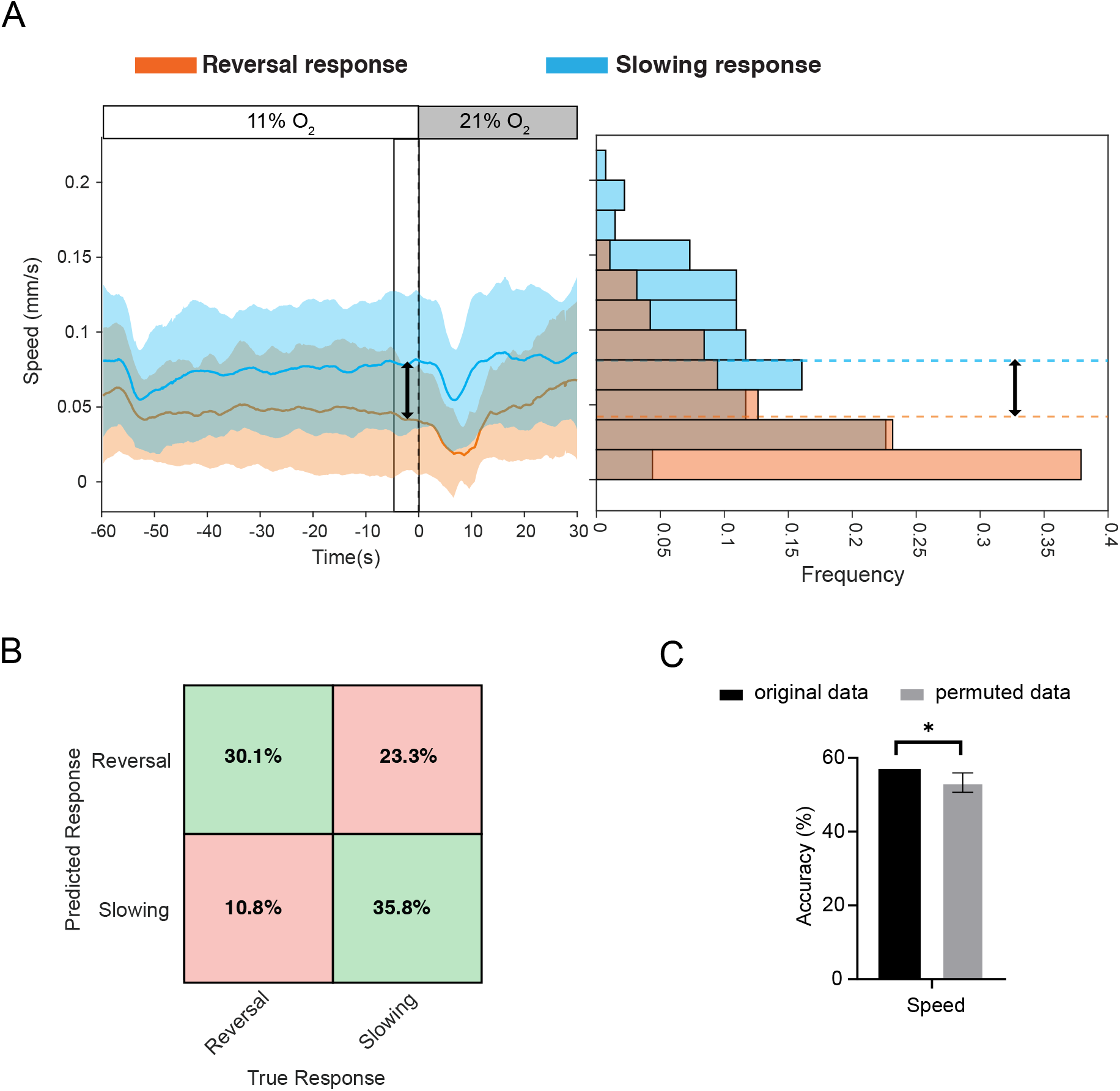
Intact predictability of speed in B-MNs:HisCl worms. Related to. Figure 2**. A.** Stimulus-triggered average forward speed of BMNs:HisCl worms in trials from two response categories (trial mean± std). The black line indicates the pre-stimulus time window (5 seconds) used to calculate the average speed values for the histogram on the right, and dotted lines show means. **B.** LDA confusion matrix showing the prediction accuracy of pre-stimulus forward speed as a predictor. Green = correctly predicted, Red = incorrectly predicted trials. **C.** Cross-validated mean LDA accuracy with actual data (Black) and shuffled data (Gray). Error bar indicates std (Permutation test, p-value *<0.05) (see methods) (n=232 trials/ N=16 animals).

**Figure S4:**
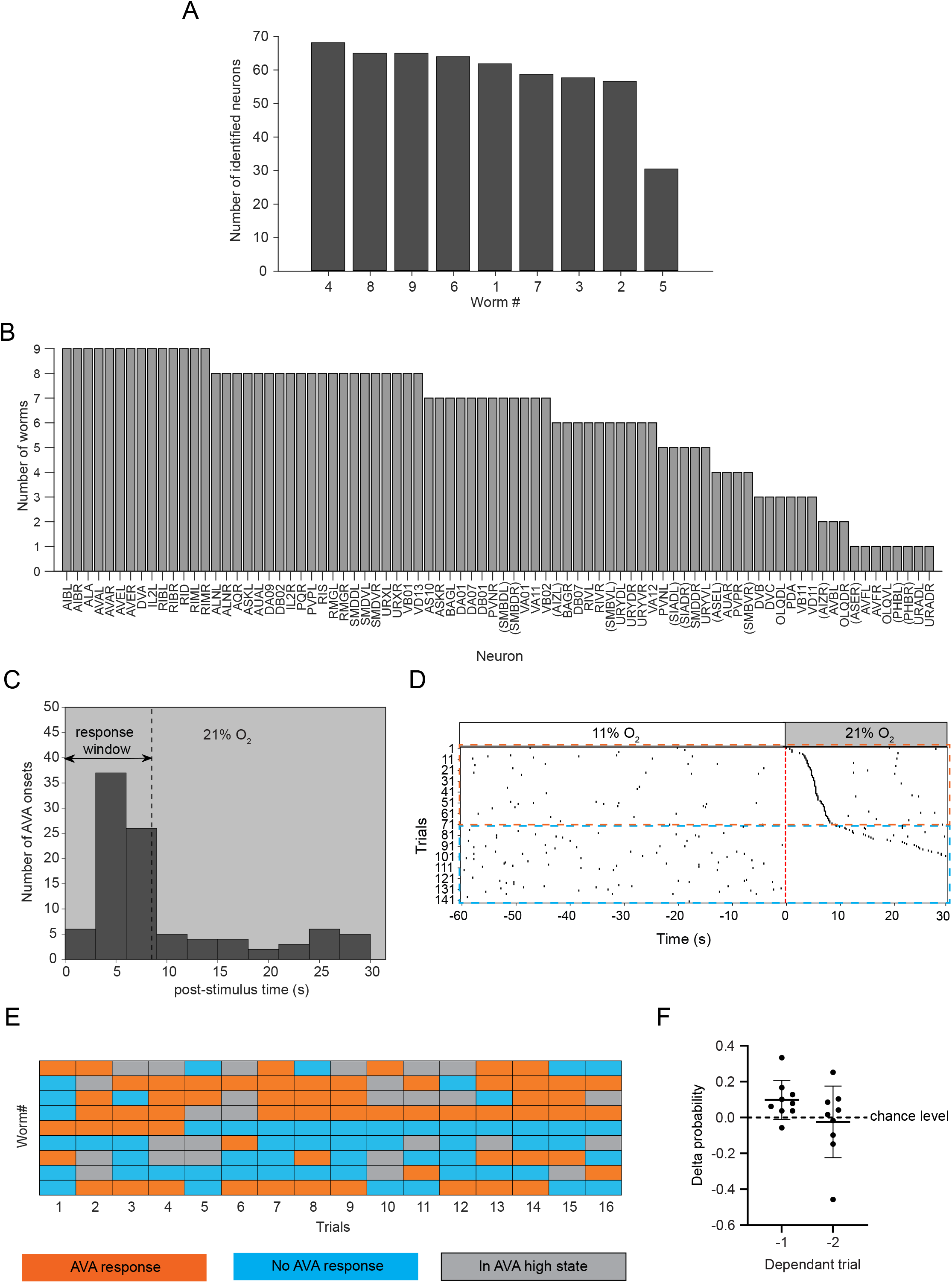
Recapitulating response variability in the immobilized imaging condition. Related to. Figures 3 **and 4. A.** Number of total neurons identified in each worm (dataset). **B.** Number of worms in which each neuron type is identified. Neurons in parentheses () indicate that neuron ID not verified and uncertain. **C.** Histogram of the first AVA rise onset time in the 21% O2 (post-stimulus) period. The black dotted line shows the defined response window (0-8.5 seconds). **D.** Stimulus-triggered occurrence of AVA rise events in all trials, sorted based on the reversal response delay within the 21% O2 stimulus period. Orange = AVA response trials and Blue = No AVA response trials, N= 9 / 117 (worms/trials). **E.** Response category of each trial. Orange = AVA/ reversal response, Blue = No AVA/slowing response, Gray = In AVA high state/already reversing. **F.** Probability of the same response in consecutive (-1) and the trial before that (-2). Each dot represents an animal, and the y-axis shows the difference from its shuffle control.

**Figure S5:**
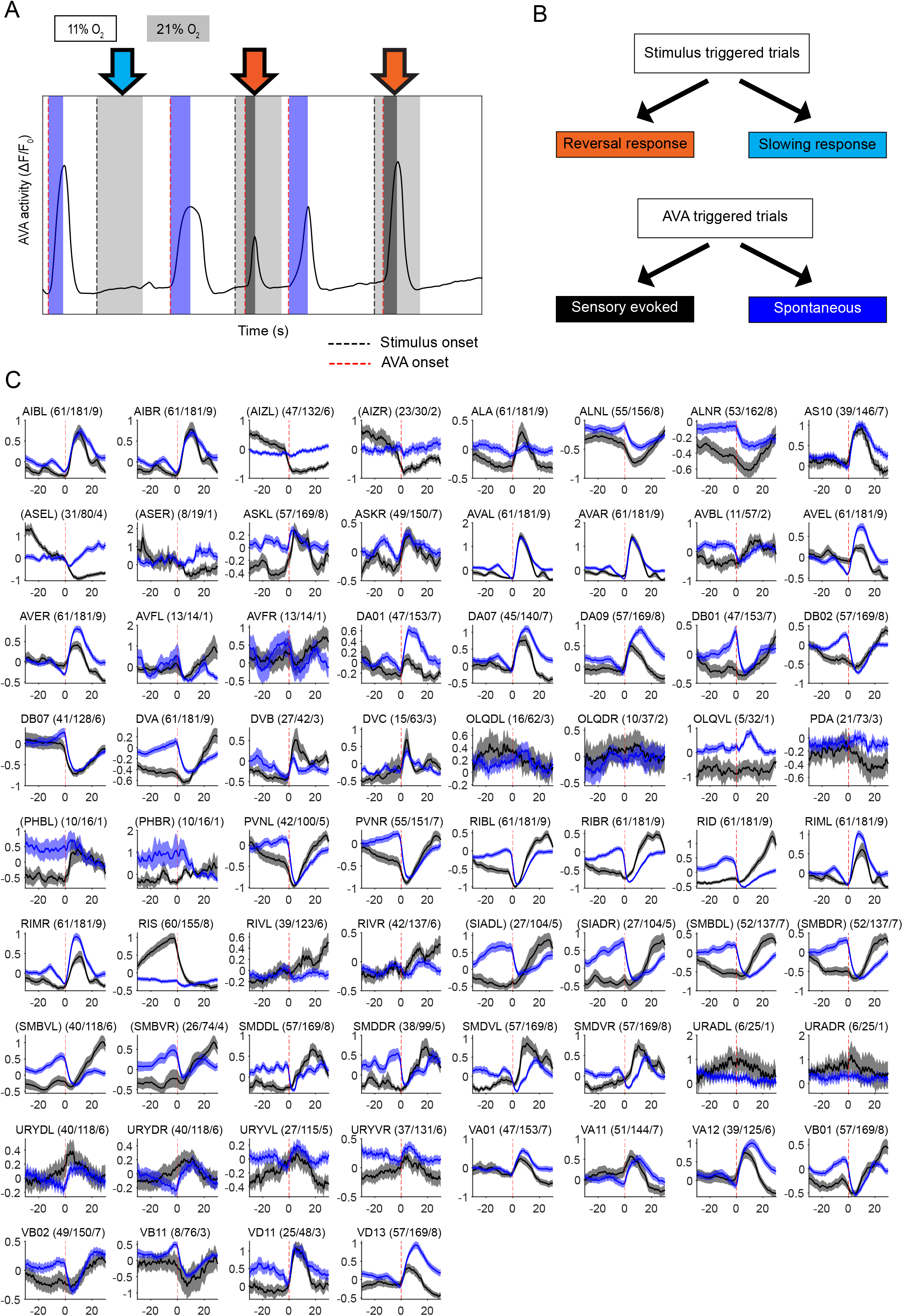
Different classifications and spontaneous vs sensory-evoked dynamics. Related to. Figure 4**. A.** Examples of spontaneous (blue) and sensory-evoked (black) reversals and reversal (orange) and slowing (cyan) trial classification based on the AVA activity trace. **B.** Diagram explaining the classifications of the data, based on AVA traces, as shown in the example in Figure S4A. **C.** Grid plot of spontaneous and sensory-evoked reversal-triggered average traces of inter-and motor neurons (mean±SEM). Each grid title contains the following: neuron name (no. of sensory-evoked reversal-triggered traces/ no. of spontaneous reversal-triggered traces/N of animals). Neurons in parentheses () indicate that neuron ID not verified and uncertain.

**Figure S6:**
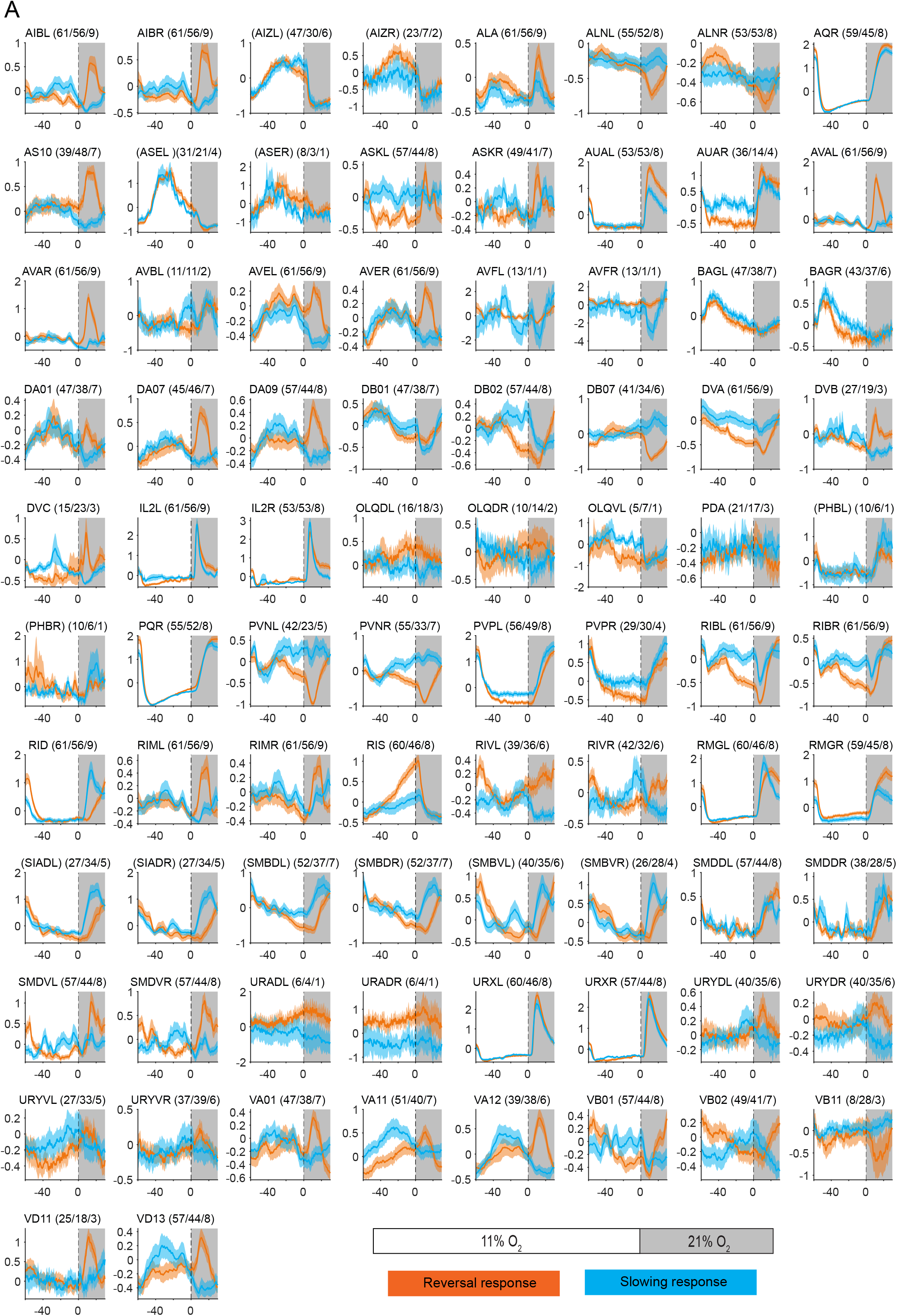
Individual neurons in reversal vs slowing response trials. Related to. Figure 5**. A.** Grid plot of stimulus-triggered average traces of all neurons in reversal and slowing response trials (mean±SEM). Each grid title contains the following: neuron name (no. of reversal response trial traces/ no. of slowing response trial traces/N of animals). Neurons in parentheses () indicate that neuron ID not verified and uncertain.

**Figure S7:**
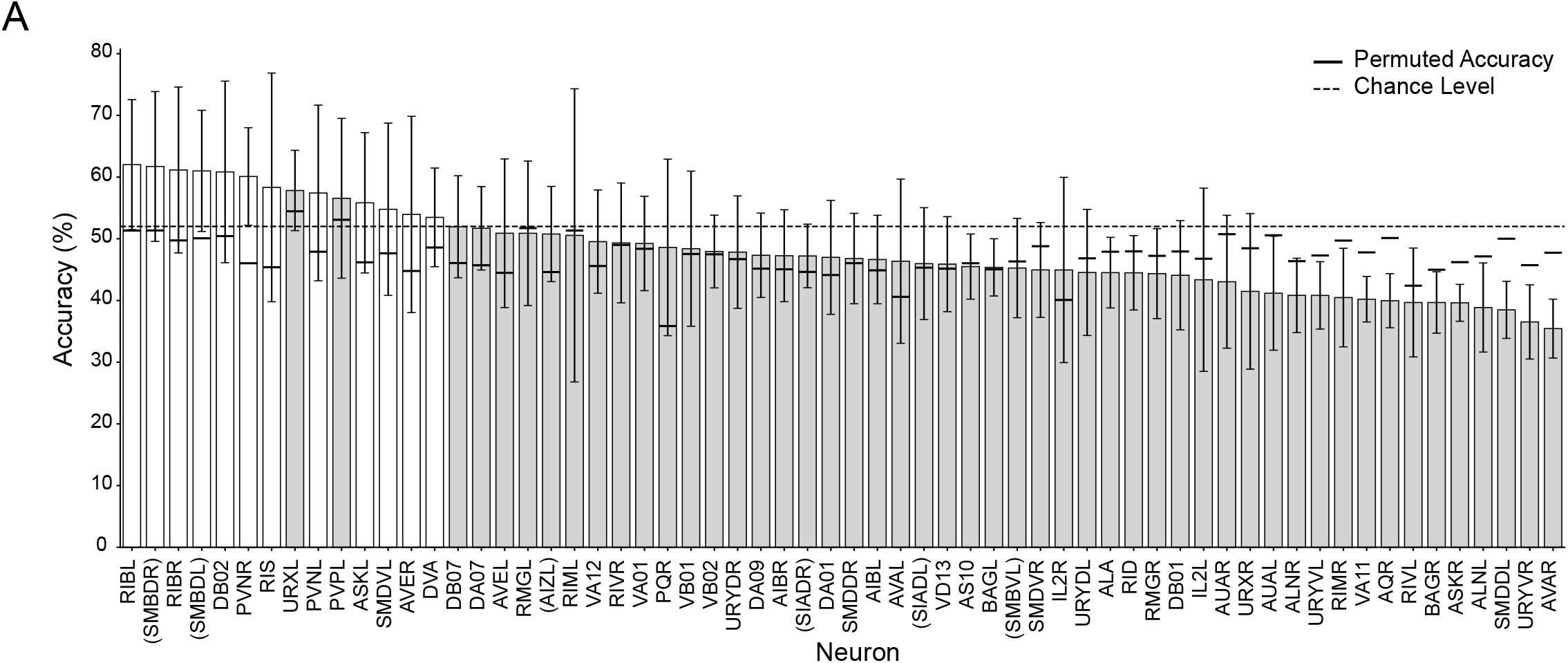
Single neuron LDA analysis. Related to. Figure 6**. A.** LDA accuracies of individual neurons with actual data (Gray and White bars, cross-validated mean± std) and shuffled data (Black line on each bar), sorted by the prediction accuracy. The dotted line indicates the chance-level prediction accuracy (∼52%). A neuron is considered significant (White bars) only when the prediction accuracy with actual data is higher than chance level, and that of shuffled data is lower than or equal to chance level (n=117 trials/ N=9 animals). Neurons in parentheses () indicate that neuron ID not verified and uncertain.

**Figure S8:**
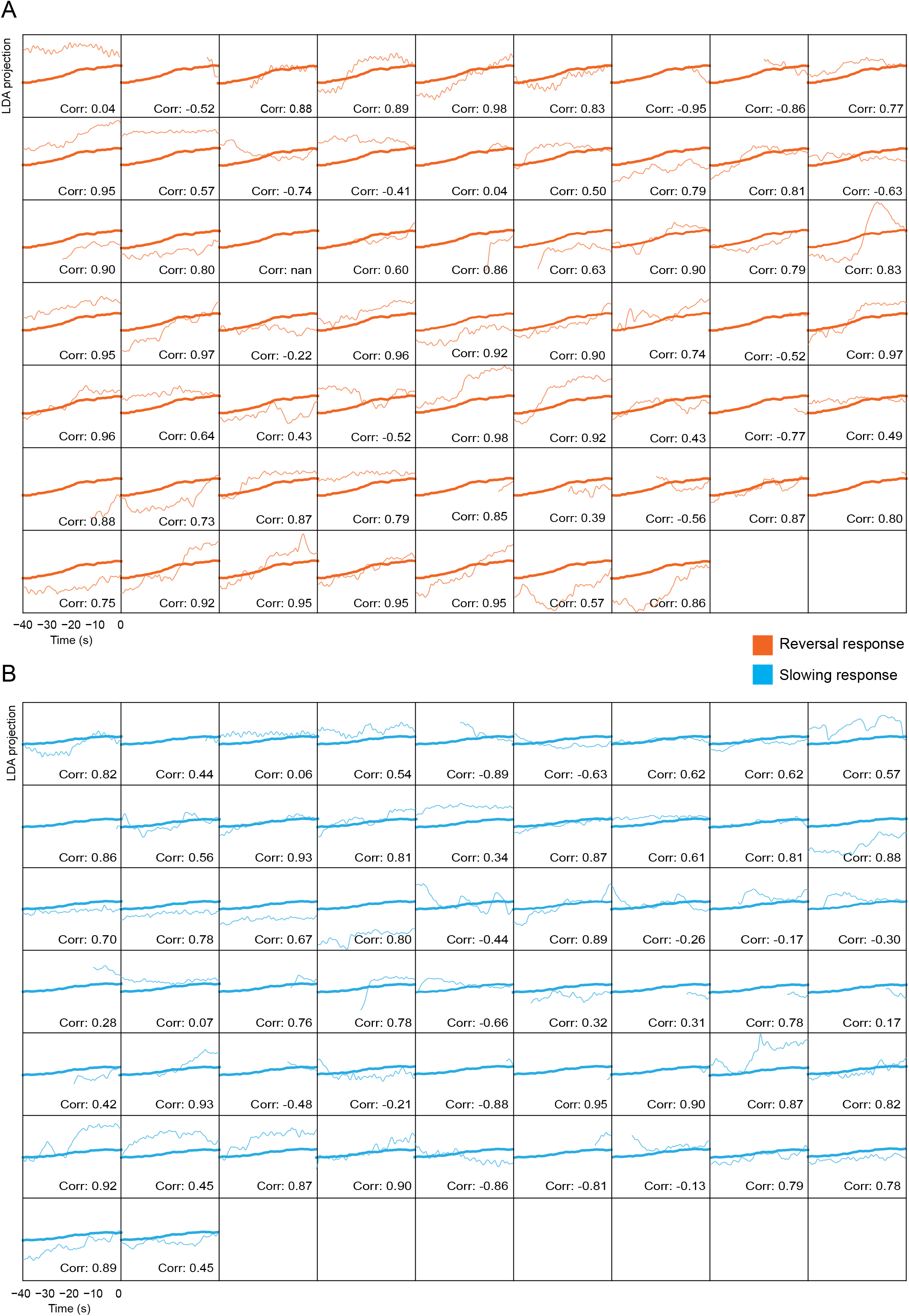
Trajectories of the LDA projection (decision variable) in individual trials. Related to. Figure 6D**. A-B.** Pre-stimulus forward state LDA projections of the PLS neuronal population model in individual reversal response trials compared to the average (bold) of all reversal response trials (A, Orange) and slowing response trials (B, Cyan). Corr: Pearson correlation coefficient.

**Figure S9:**
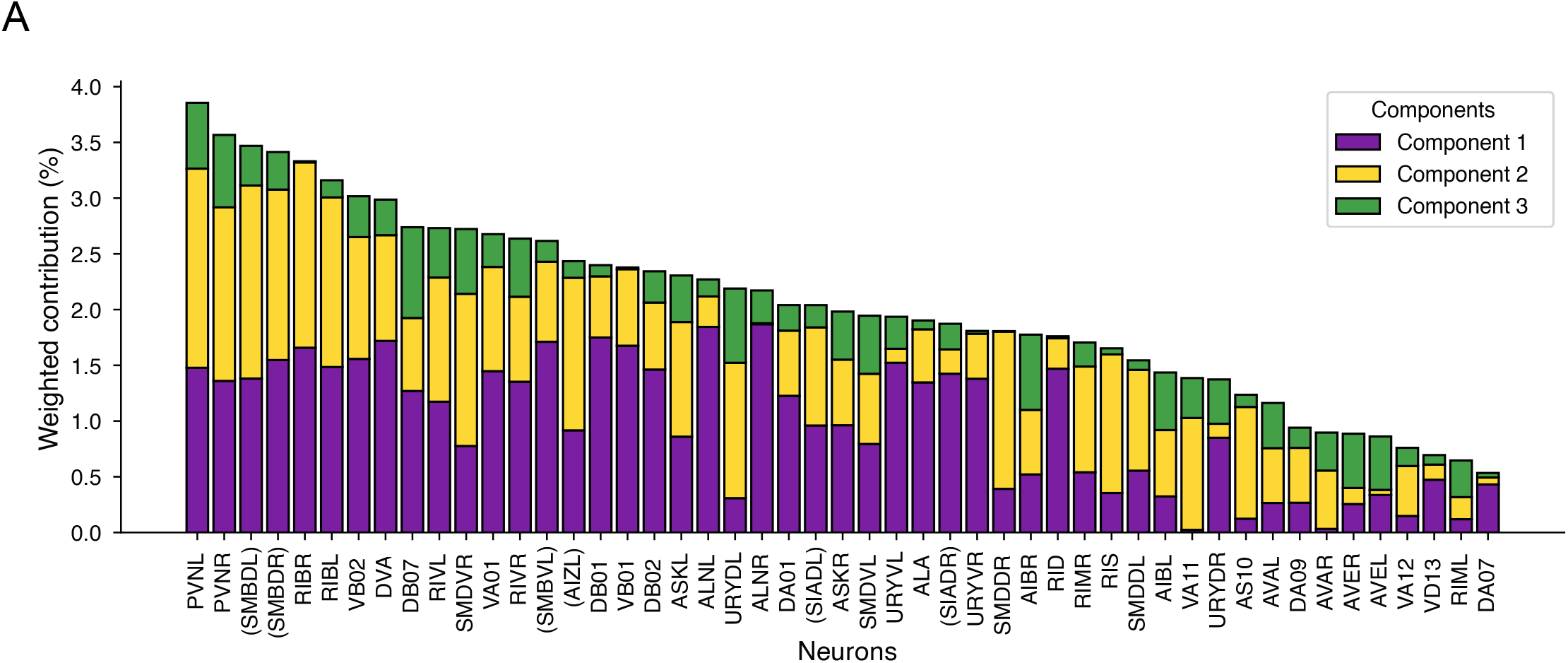
A. Weighted contribution (loadings) of each neuron to the top 3 PLS dimensions (components), indicating individual neurons’ contributions (feature importance) to the spontaneous PLS model. Feature importance of each component (see methods). Neurons in parentheses () indicate that neuron ID not verified and uncertain.

**Figure S10:**
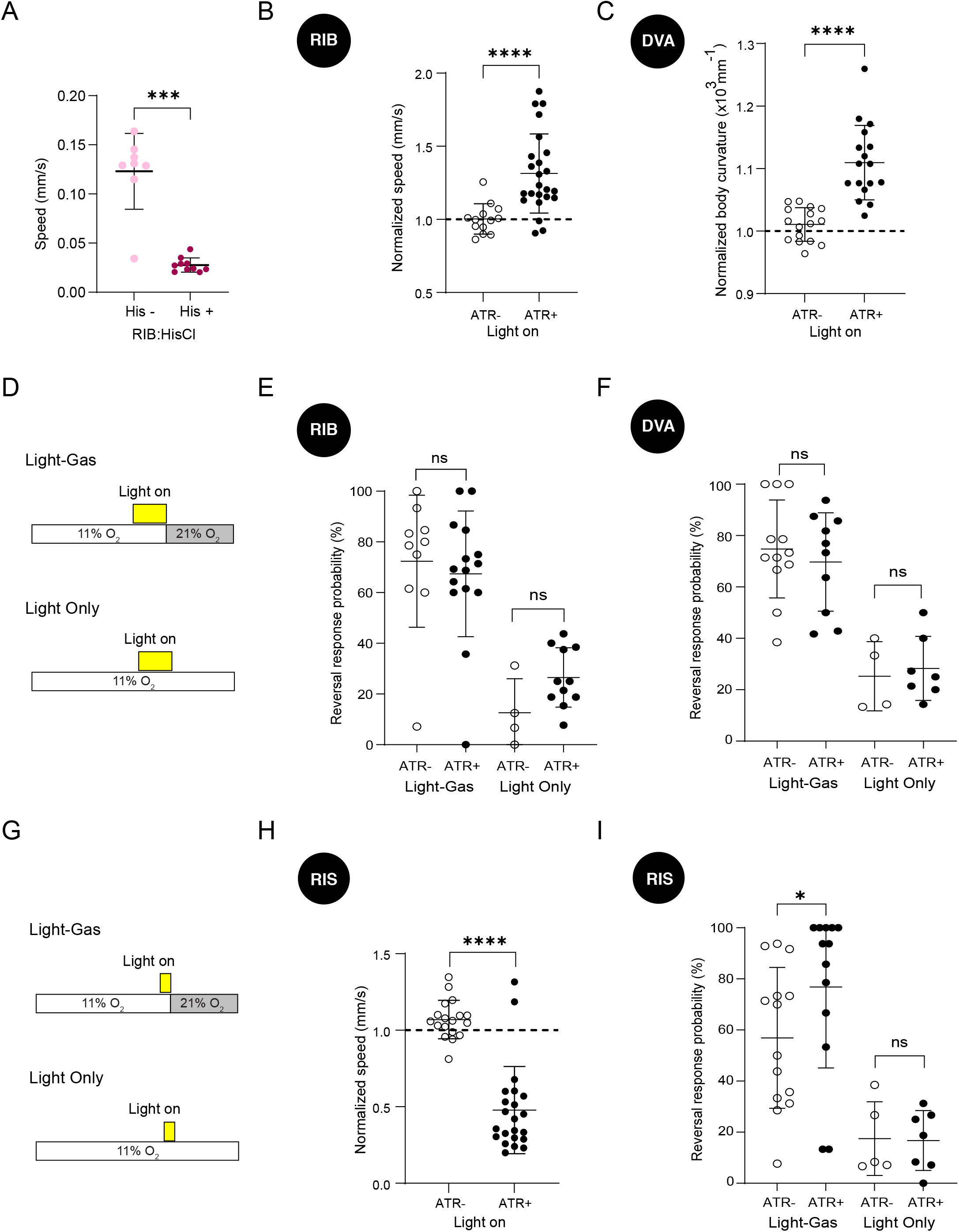
Experimental data supporting neuron perturbation experiments. Related to. Figure 8**. A.** Average baseline (11% O2) speed of RIB:HisCl worms. **B.** Average fold change in speed (mean speed in light-on period divided by the mean speed in light-off period in that trial) across trials in RIB Chrimson worms. **C.** Average fold change in curvature (mean curvature in light-on period divided by the mean curvature in light-off period in that trial) across trials in DVA Chrimson worms. **D.** Schematic of experimental design for RIB and DVA optogenetic experiments coupled with gas stimulus. Reversal probability of **E.** RIB and **F.** DVA Chrimson worms in the light-gas coupled stimulation (experiment) and light-only stimulation (control) conditions. The mean of each control conditions was used for calculating the corresponding normalized reversal probabilities shown in Fig. 8D & E (see methods). **G.** Schematic of experimental design for RIS optogenetic experiments. **H.** Average fold change in speed (mean speed in light-on period divided by the mean speed in light-off period in that trial) across trials in RIS Chrimson worms. **I.** Reversal probability of RIS Chrimson worms in the light-gas coupled stimulation (experiment) and light-only stimulation (control) conditions. The mean of each control condition was used for calculating the corresponding normalized reversal probabilities shown in Fig. 8F. Each dot represents an animal. The error bar shows mean± std (Mann-Whitney test p-value ****<0.0001, ***<0.001, **<0.01, *<0.05).

**Table S1:** Strain list.

| Strain name | Genotype | Description | Relevant Figures | Reference |
| --- | --- | --- | --- | --- |
| AX1858 | *npr-1(ad609); lite-1(ce314)* |  | Fig. 2 & 8 | (J. Liu et al. 2010; Bono and Bargmann 1998) |
| ZIM2344 | *mzmls52; mzmls120; npr-1(ad609); lite-1(ce314)* | *mzmls52* = [*mzmEx771* = [*unc-31::NLSGCaMP6f*]<br>*mzmls120* = [*mzmEx1341* = [*mlc-2::HisCl::rpl-3 3'UTR; flp-17::scarlet*] | All Figures | (El Mouridi et al. 2021; Kaplan et al. 2020; Stefanakis et al. 2015) |
| ZIM2503 | *mzmEx1365; npr-1(ad609); lite-1(ce314)* | *mzmEx1365* = [*unc-129(DB)::HisCl::SL2::mCherry; ceh-12short::HisCl1::SL2::mCherry; unc-122::GFP*] | Fig. 2 | (Gao et al. 2018) |
| ZIM2461 | *mzmEx1333; npr-1(ad609); lite-1(ce314)* | *mzmEx1333* = [*sto-3::HisCl::mCherry; flp-17::mCherry*] | Fig. 8 | (Kaplan et al. 2020) |
| ZIM2478 | *mzmEx928; npr-1(ad609); lite-1(ce314)* | *mzmEx928* = [*sto-3::Chrimson::SL2::mCherry; unc-122::GFP*] | Fig. 8 | (Kaplan et al. 2020) |
| ZIM2500 | *mzmEx1107; mzmEx1083; npr-1(ad609); lite-1(ce314)* | *mzmEx1107* = [*aptf-1::CRE::aptf-1 3'UTR; flp-17::cherry*], *mzmEx1083* = [*flp-11::DIO-HisCl::SL2::mCherry; unc-122::dsRed*] | Fig. 8 | (Turek et al. 2016) |
| ZIM1878 | *mzmEx1107; mzmEx1084; npr-1(ad609); lite-1(ce314)* | *mzmEx1107* = [*aptf-1:CRE:aptf-1 3'UTR; flp-17::cherry*], *mzmEx1084* = [*flp-11::DIO-ChrimsonSL2mCherry; unc-122::dsRed*] | Fig. 8 | (Turek et al. 2016) |
| ZIM2523 | *mzmEx1362; npr-1(ad609); lite-1(ce314)* | *mzmEx1362* = [*nlp-12::HisCl-SL2-mCherry; flp-17::mScarlet*] | Fig. 8 | (Hums et al. 2016) |
| ZIM2517 | *mzmEx1374; npr-1(ad609); lite-1(ce314)* | *mzmEx1374* = [*nlp-12::Chrimson::SL2::mCherry; unc-122::GFP*] | Fig. 8 | (Hums et al. 2016) |
| ZIM2376 | *mzmls120; otIs669; Mzmls52; npr-1(ad609); lite-1(ce314)* | *mzmls120* = [*mzmEx1341* = [*mlc-2::HisCl::rpl-3 3'UTR; flp-17::scarlet*], *otIs669* [NeuroPAL] V, *Mzmls52* = [*mzmEx771* = [*unc-31::NLSGCaMP6f*] | Fig. S4B | (Yemini et al. 2021) |

**Table S2:** Modulation index of individual neurons Neurons in parenthesis () indicate that neuron ID not verified and uncertain.

| Neuron name | Modulation index |
| --- | --- |
| URXL | 0.87 |
| URXR | 0.85 |
| IL2R | 0.84 |
| IL2L | 0.83 |
| AUAL | 0.79 |
| RMGL | 0.79 |
| RMGR | 0.78 |
| AQR | 0.71 |
| RIS | 0.68 |
| PQR | 0.68 |
| AUAR | 0.62 |
| PVPL | 0.61 |
| PVPR | 0.57 |
| ALA | 0.50 |
| (AIZL) | 0.47 |
| BAGR | 0.43 |
| (SIADL) | 0.34 |
| URADR | 0.33 |
| BAGL | 0.29 |
| RID | 0.29 |
| (SMBVR) | 0.29 |
| (AIZR) | 0.28 |
| URADL | 0.21 |
| DVB | 0.19 |
| ASKL | 0.18 |
| OLQDR | 0.17 |
| AVBL | 0.13 |
| SMDDR | 0.11 |
| (SMBVL) | 0.11 |
| URYDL | 0.09 |
| (SIADR) | 0.09 |
| ASKR | 0.09 |
| SMDDL | 0.08 |
| RIVL | 0.07 |
| SMDVL | 0.07 |
| (SMBDL) | 0.07 |
| RIBL | 0.05 |
| URYVL | 0.05 |
| RIBR | 0.04 |
| URYVR | 0.03 |
| PDA | 0.01 |
| DVA | 0.01 |
| SMDVR | -0.01 |
| (SMBDR) | -0.02 |
| DB01 | -0.03 |
| PVNL | -0.03 |
| RIVR | -0.05 |
| OLQVL | -0.06 |
| (ASEL) | -0.07 |
| VB01 | -0.08 |
| (ASER) | -0.10 |
| PVNR | -0.11 |
| VD11 | -0.11 |
| VA11 | -0.15 |
| ALNR | -0.17 |
| OLQDL | -0.19 |
| DB02 | -0.25 |
| AIBR | -0.27 |
| AVAL | -0.28 |
| AVAR | -0.28 |
| URYDR | -0.29 |
| ALNL | -0.30 |
| VA01 | -0.34 |
| AIBL | -0.35 |
| DB07 | -0.39 |
| VA12 | -0.39 |
| RIMR | -0.40 |
| DVC | -0.41 |
| DA09 | -0.42 |
| DA07 | -0.45 |
| RIML | -0.47 |
| VD13 | -0.48 |
| VB11 | -0.48 |
| AS10 | -0.48 |
| AVER | -0.52 |
| AVEL | -0.58 |
| VB02 | -0.60 |
| DA01 | -0.68 |

## Notes

### Competing Interest Statement

The authors have declared no competing interest.

